# The *Arabidopsis* SR45 splicing factor bridges the splicing machinery and the exon-exon junction complex

**DOI:** 10.1101/2023.08.07.552250

**Authors:** Steven Fanara, Marie Schloesser, Marine Joris, Simona De Franco, Marylène Vandevenne, Frédéric Kerff, Marc Hanikenne, Patrick Motte

**Affiliations:** InBioS-PhytoSystems, Functional Genomics and Plant Molecular Imaging, University of Liège, 4000, Liège, Belgium; InBioS-Center for Protein Engineering, Laboratory of Biological Macromolecules, University of Liège, 4000, Liège, Belgium; InBioS-Center for Protein Engineering, Laboratory of Crystallography, University of Liège, 4000, Liège, Belgium; InBioS-PhytoSystems, Translational Plant Biology, University of Liège, 4000, Liège, Belgium

**Keywords:** Serine/arginine-rich splicing factor, SR45, Arabidopsis, splicing, exon-exon junction complex, EJC, protein-RNA and protein-protein interaction, ASAP, PSAP

## Abstract

The Arabidopsis splicing factor serine/arginine-rich 45 (SR45) contributes to several biological processes. The *sr45-1* loss-of-function mutant exhibits delayed root development, late flowering, unusual numbers of floral organs, shorter siliques with decreased seed sets, narrower leaves and petals, and altered metal distribution. SR45 bears a unique RNA recognition motif (RRM) flanked by one serine/arginine-rich (RS) domain on both sides. Here, we studied the function of each of SR45 domains by examining their involvement in: (i) the spatial distribution of SR45, (ii) the establishment of a protein-protein interaction network including spliceosomal and exon-exon junction complex (EJC) components, and (iii) the RNA binding specificity. We report that the endogenous *SR45* promoter is active during vegetative and reproductive growth, and that the SR45 protein localizes in the nucleus. We demonstrate that the C-terminal arginine/serine-rich domain is a determinant of nuclear localization. We show that the SR45 RNA recognition motif (RRM) domain specifically binds purine-rich RNA motifs via three residues (H101, H141, Y143), and is also involved in protein-protein interactions. We further show that SR45 bridges both mRNA splicing and surveillance machineries as a partner of EJC core components and peripheral factors, which requires phosphoresidues likely phosphorylated by kinases from both CLK and SRPK families. Our findings provide insights into the contribution of each SR45 domain to both spliceosome and EJC assemblies.

**Highlight:** The contribution of the Arabidopsis SR45 splicing factor individual domains to its nuclear localization, ability to contact *in planta* novel protein partners and specifically bind RNA motifs was examined.

## Introduction

The precise recognition of splice sites at exon-intron boundaries is crucial in order to accurately perform precursor-messenger RNA (pre-mRNA) splicing (Wahl *et al*., 2009). While constitutive splicing results from the exclusion of each intron from the pre-mRNA, alternative splicing (AS) can lead to intron retention, shortened or lengthened exon or the skipping of an entire exon (Reddy *et al*., 2013; Wang *et al*., 2015). This occurs when weak or non-canonical splice sites are recognized and processed by the splicing machinery (Chen and Manley, 2009). It has important consequences for mRNA abundance and protein stability (Graveley, 2001). AS can be linked to an mRNA degrading process called nonsense-mediated mRNA decay (NMD) through the emergence of premature termination codons (PTCs) in alternatively spliced transcripts, which ultimately regulates transcript levels (Palusa and Reddy, 2010; Kalyna *et al*., 2012). Protein isoforms that arise from alternative splicing usually differ in their primary sequence and domain organization which often alters their localization, stability, function or ability to interact with their protein or RNA partners (Laloum *et al*., 2018).

Serine/arginine-rich (SR) proteins constitute a highly conserved family in multicellular eukaryotes (Califice *et al*., 2012), sharing a modular structure characterized by the presence of one or two RNA recognition motif(s) [RRM(s)] in the N-terminus and a C-terminal domain rich in arginine-serine dipeptide repeats (RS) (Manley and Krainer, 2010; Califice *et al*., 2012). While the RNA binding affinity is mainly dictated by the RRM domain (Maris *et al*., 2005; Daubner *et al*., 2013), the RS domain is involved in protein-RNA and protein-protein interactions through reversible phosphorylation (Xiao and Manley, 1997; Shen *et al*., 2004; Schütze *et al*., 2016; Schwich *et al*., 2021). The phosphorylation status of the RS domain is controlled by at least two conserved protein kinase families, serine arginine protein kinases (SRPK) and *cdc2*-like kinases (CLK), as wells as some phosphatases (e.g., <u>P</u>rotein <u>P</u>hosphatase <u>1</u>, PP1) (Aubol *et al*., 2016; Stankovic *et al*., 2016; Aubol *et al*., 2017). The SRPK kinases are located in the cytoplasm and the nucleus and strictly phosphorylate serine-arginine dipeptides (Ma *et al*., 2009; Koutroumani *et al*., 2017). They influence the localization of SR proteins by supporting their phosphorylation-dependent transport into the nucleus via an SR-specific transportin protein (TRN-SR, TNPO3) (Kataoka *et al*., 1999; Lai *et al*., 2000; Lai *et al*., 2001; Xu *et al*., 2011). The CLK members are dual kinases and strictly nuclear (Lee *et al*., 1996). They phosphorylate not only serine-arginine but also serine-proline dipeptides (Lee *et al*., 1996; Aubol *et al*., 2013). Some SR proteins display a nucleocytoplasmic dynamics and are known to play many additional roles in (pre-)mRNA processing in mammals, including transcription, mRNA export (Huang and Steitz, 2001), stability and surveillance (Zhang and Krainer, 2004), and translation (Sanford *et al*., 2004). Based on the sole phylogeny of the RRM, the *Arabidopsis thaliana* (Arabidopsis) genome possesses nineteen SR protein-encoding genes divided into seven subgroups, four of which are plant-specific because of a unique multidomain structure (RS, SCL, RS2Z and SR45 subfamilies) (Barta *et al*., 2010; Califice *et al*., 2012).

SR45 is an atypical SR protein because it possesses a unique RRM flanked by one RS domain on each side (Barta *et al*., 2010), a domain organization similar to the one of the human splicing factors <u>Tra</u>nsformer-<u>2β</u> (Tra2β) and RNPS1 (<u>RN</u>A-binding <u>P</u>rotein with <u>S</u>er-rich domain <u>1</u>) (Cléry *et al*., 2011), the human ortholog of SR45 (Califice *et al*., 2012). RNPS1 plays critical roles in splicing (Robinson *et al*., 1999) and NMD by coupling the deposition of a quality control complex (<u>E</u>xon <u>J</u>unction <u>C</u>omplex, EJC) on spliced mRNAs to the <u>UP</u>-<u>F</u>RAMESHIFT (UPF) protein complex, which consists of UPF1, UPF2 and UPF3 and triggers NMD (Lykke-Andersen *et al*., 2001). Two scaffold proteins, ACINUS and PININ, form with the EJC peripheral factors RNPS1 and SAP18 (<u>S</u>in3A <u>A</u>ssociated <u>P</u>rotein <u>18</u>) the ternary complexes ASAP and PSAP, respectively, in order to regulate AS and NMD (Sakashita *et al*., 2004; Singh *et al*., 2010; Murachelli *et al*., 2012; Schlautmann *et al*., 2022). The EJC itself is a conserved protein complex comprising four highly conserved core proteins named <u>e</u>ukaryotic initiation factor <u>4A3</u> (eIF4AIII), Mago nashi (MAGOH/Mago), Y14 and <u>M</u>etastatic <u>L</u>ymph <u>N</u>ode <u>51</u> (MLN51) (Tange *et al*., 2005). The cytoplasmic disassembly of the EJC results from the action of PYM (Bono *et al*., 2004). This EJC core serves as a platform for multiple peripheral proteins in order to coordinate several steps of mRNA processing, namely (alternative) splicing, intra-cellular RNA localization (which notably involves ALY/REF), translation efficiency and mRNA stability control by NMD (Le Hir *et al*., 2001; Nott *et al*., 2004; Singh *et al*., 2012).

In Arabidopsis, little is known about the functional characteristics of the EJC components and the associated peripheral factors (Chen *et al*., 2019). The Arabidopsis homologues of MAGO, Y14, PYM and eIF4AIII have been previously described (Johnson *et al*., 2004; Park and Muench, 2007; Park *et al*., 2009). MAGO and Y14 can heterodimerize (Park and Muench, 2007). While the C-terminal portion of MAGO is critical for this interaction, it is dispensable for the association with eIF4AIII (Cilano *et al*., 2016). PYM interacts with the MAGO-Y14 complex, as well as with the isolated proteins (Park and Muench, 2007). PYM and Y14 associate only when both proteins are dephosphorylated, but the phosphorylation state does not affect the association with MAGO (Mufarrege *et al*., 2011). Knockdown of *MAGO* in plants leads to several developmental defects, including vegetative and reproductive development impairment (Park *et al*., 2009). The DEAD-box RNA helicase eIF4AIII is a nuclear protein whose subcellular localization is altered under hypoxia (Koroleva *et al*., 2009). It interacts with ALY4 *in vitro* and colocalizes with MAGO and Y14 *in planta* (Koroleva *et al*., 2009). While RNPS1 was shown to interact with Y14, no link was established between SR45 and Y14 or any other EJC core component in Arabidopsis (Shi and Xu, 2003; Hsu Ia *et al*., 2005; Wang *et al*., 2018). However, it was recently shown that ASAP and PSAP complexes containing SR45 exist in Arabidopsis and contribute to the regulation of *FLC* expression (Qüesta *et al*., 2016; Chen *et al*., 2019; Bi *et al*., 2021; Mikulski *et al*., 2022) and to the reproductive transition (Bi *et al*., 2021). SR45 was shown to affect the accumulation of ASAP protein components (Chen *et al*., 2019).

As SR45 presents an atypical modular structure among SR proteins, with a unique RRM flanked by an unstructured RS domain on both sides **(Supplementary Figure S1)**, the function of these domains in the modulation of the SR45 splicing activity, i.e. its localization, ability to interact with proteins and specificity of RNA binding, was examined. We first studied the expression profile of *SR45* during plant development with native promoter-driven reporter constructs to link previously described phenotypes of the loss-of-function mutant to the tissues where SR45 is expressed. By substituting serines from both RS domains to alanines, we established that the SR45 RS domains are required to retain the nuclear localization and play a key role in protein-protein interactions. We further determined the RNA motif recognized by SR45 and demonstrated how three amino acids in the RRM define the RNA binding specificity. Finally, we established how phosphorylatable serines are critical for establishing the SR45 protein-protein interaction network, particularly with kinases, EJC core components and EJC peripheral factors. Our findings provide new insights into the involvement of SR45 in the recruitment of several splicing factors and in EJC assembly.

## Materials and methods

### Plant material and growth conditions

Arabidopsis plants (Col-0 ecotype) were stably transformed by floral dipping, and independent T3 homozygous lines were analyzed. All observations were performed in at least three independent lines. Seeds were surface-sterilized, then germinated on 1/2 Murashige and Skoog (MS) medium (Duchefa Biochimie) supplemented with sucrose (1% w/v, Duchefa Biochimie) and Select Agar (0.8% w/v, Sigma-Aldrich) after stratification in the dark at 4°C for 48h. For expression profiling, 3-week-old seedlings were grown in hydroponic trays (Araponics) in Hoagland medium for ten weeks (until silique development) as previously reported (Talke *et al*., 2006; Hanikenne *et al*., 2008). Hydroponics were grown under a 16-h light (100 μmol m^-2^ sec^-1^)/8-h dark regime in a climate-controlled growth chamber (21°C). The nutrient medium was renewed with fresh solution once a week.

*Nicotiana tabacum* (cv Petit Havana) transient transformations by *Agrobacterium* infiltration were performed as described (Rausin *et al*., 2010; Stankovic *et al*., 2016).

### Vector constructions

All binary vector constructions and primers used in this study are listed and detailed in **Supplementary Tables S1-9**. All PCRs were carried out using the Q5 DNA polymerase (NEB) on Arabidopsis cDNA libraries or genomic DNA. All constructs were verified by sequencing. For plant transient and stable transformations, all final plasmids were electroporated into the *Agrobacterium tumefaciens* strain GV3101 (pMP90) and subsequently used for agroinfiltration or floral dipping, respectively. All constructions contain the coding sequence of the *SR45.1* isoform, which we name *SR45* hereafter (Zhang and Mount, 2009).

For yeast two- or three-hybrid assays (Y2H or Y3H), cDNAs coding for the potential interactors were cloned into pGBKT7 and pGADT7 vectors (Clontech) using different restriction sites, with the exception of the *SR45* and mutant variant coding sequences that were cloned into pGBKT7(+) and pGADT7(+) in which *Asc*I and *Pac*I restriction sites were added (see **Supplementary Tables S1 and S2** for primer lists).

The promoter region of *SR45* [1612 bp upstream of the ATG, as described by (Zhang and Mount, 2009)] was amplified by PCR from genomic DNA. The promoter amplicon was ligated at the *Sbf*I/*Kpn*I sites of the pMDC100 vector (Curtis and Grossniklaus, 2003) to create the pMDC100:pSR45 vector. The *SR45* open reading frame (1242 bp, from the ATG to the last codon before the stop codon) was then cloned at *Asc*I/*Pac*I sites to obtain pMDC100:pSR45:SR45 in which a *nos* terminator (tNOS) was added between *Sac*I restriction sites. Finally, the EGFP coding sequence was cloned at the *Pac*I site to create the pMDC100:pSR45:SR45:EGFP:tNOS vector. To construct the pMDC32:pSR45:EGFP vector, the 2X35S promoter of the pMDC32 vector (Curtis and Grossniklaus, 2003) was replaced by the *SR45* promoter and EGFP was cloned downstream of the promoter at *Kpn*I and *Sac*I restriction sites **(Supplementary Table S3)**.

The *SR45mutRRM* version was obtained by PCR-based site-directed mutagenesis as described (Rausin *et al*., 2010; Stankovic *et al*., 2016) **(Supplementary Table S4)**. The other mutated *SR45* versions were assembled by PCR from partially overlapping amplicons, which corresponded to the coding sequence of each domain of SR45. They were amplified using either the native *SR45* gene, the *SR45mutRRM* gene, or the two RS mutated domains (named RA1 and RA2) synthesized by Eurofins Genomics (see **Supplementary Tables S5 and S6**). All assembled mutant versions were then introduced into yeast vectors for Y2H (at *Asc*I/*Bam*HI) or pMDC32 fused to EGFP for protein localization (at *Asc*I/*Pac*I sites), respectively.

The BiFC constructs were obtained as described (Stankovic *et al*., 2016) (see **Supplementary Tables S7** for primer lists). The TriFC was performed with an additional plasmid, pMDC32 (Curtis and Grossniklaus, 2003), containing no coding sequence or the one of either the RRM domain of ACINUS (765 bp corresponding to the RRM domain and its C-terminal extension, from the 1141^st^ base to the stop codon and preceded by an ATG codon) or PININ.

For SELEX experiments, sequences encoding the RRM of SR45, and the RRM1 or the RRM2 of SR34 were cloned at *Bam*HI/*Eco*RI sites of the pGEX6P-1 as described (De Franco *et al*., 2019) **(Supplementary Table S8)**. Sequences encoding mutated versions of RRM1 and RRM2 of SR34 were amplified from vectors previously described (Stankovic *et al*., 2016).

### Yeast two- and three-hybrid systems

The vectors and yeast (*Saccharomyces cerevisiae*) strains provided in the Matchmaker Gold Yeast Two-Hybrid System (Clontech) were used to perform directed interaction analyses (dY2H). Prior to (directed) Y2H experiments, toxicity and autoactivation of the wild-type *SR45* and mutant variants were tested simultaneously on a medium -Trp/X-a-gal [for pGBKT7/pGBKT7(+)] and -Leu/X-a-gal [for pGADT7/pGADT7(+)] using 100 ng of plasmid. Yeast growth was not affected by any of the constructs, none of which led to autoactivation of the reporter gene (*MEL1*) **(Supplementary Figure S2)**. Dimeric interactions were then tested on selective medium -Trp/-Leu/-His/X-a-gal/Aureobasidin A.

For Y2H screening, the Y2HGold strain was transformed with the bait vector (pGBKT7-SR45) and then mated with strain Y187 containing an Arabidopsis cDNA library (Mate and Plate Library-Universal Arabidopsis, Clontech) (Stankovic *et al*., 2016) or a custom cDNA library made from entire mature leaf material (Make Your Own “Mate & Plate” Library System, Clontech). Trimeric interactions (Y3H) were tested on selective medium -Met/-Trp/-Leu/-His/X-a-gal/Aureobasidin A: yeast expressing SAP18 fused to the binding domain of GAL4 (using SR45 as a bridge) were mated with yeast expressing either ACINUS or PININ fused to the activation domain of GAL4.

### Confocal Microscopy

Live cell imaging was performed on Leica TCS SP2 and SP5 inverted confocal laser microscopes (Leica Microsystems). A water-immersion objective was used to collect images at 512×512 pixel resolution. EGFP was visualized using the 488-nm line of the argon laser, and the emission light was dispersed and recorded at 500 to 560 nm (Rausin *et al*., 2010; Stankovic *et al*., 2016; Joris *et al*., 2017). For BiFC and TriFC experiments, YFP was excited at 514 nm using the argon laser and the emission light was recorded at 525 to 600 nm.

### Analysis of the mRNA levels

100 mg of plant tissues were used to extract total RNA using NucleoSpin RNA Plant kit (Macherey Nagel) as per manufacturer’s instruction. cDNAs were synthesized from 1 µg of total RNAs using the RevertAid H Minus First Strand cDNA Synthesis Kit (Fisher Scientific) with oligo(dT) primers. Quantitative PCR reactions were performed in 384-well plates using a QuantStudio5 system (Applied Biosystems) and Takyon Low Rox SYBR MasterMix dTTP Blue (Eurogentec). Samples were obtained from three independent biological experiments. A total of three technical replicates were performed for each combination of cDNA and primer pair **(Supplementary Table S9)**. Gene expression was normalized relative to At1g58050 as described (Pfaffl, 2001). *At1g58050* expression was the most stable gene among all tested references (*EF1*α and *UBQ10*) (Czechowski *et al*., 2005; Spielmann *et al*., 2020).

### Production of recombinant proteins

The recombinant protein production was performed using the expression vector pGEX6P-1 in *E. coli* BL21(DE3) cells as described (De Franco *et al*., 2019). Protein productions (native and mutated RRM of SR45, or RRM1 or RRM2 of SR34) were carried out overnight at 18°C upon induction with 0.5 mM IPTG. Lysis of the cells was carried out using a sonicator in 50LmM Tris-HCl pH 8, 1 M NaCl, 2LmM MgCl_2_, 1 mM dithiothreitol (DTT) and 1 mM PMSF. GST fusion proteins were purified on Glutathione-Sepharose^®^ 4B beads (GE Healthcare) and eluted using a solution of 50 mM Tris-HCl pH 8, 100 mM NaCl, 1LmM dithiothreitol (DTT) and 35 mM reduced glutathione.

### SELEX experiments

The Systematic Evolution of Ligands by EXponential enrichment (SELEX) experiment was performed using a randomized 25-nt DNA template library as previously reported (De Franco *et al*., 2019). 500 ng-1 µg of the purified template DNA library was used for the *in vitro* transcription reaction (T7 RiboMAX™ Express Large Scale RNA Production System, Promega). Unincorporated nucleotides were removed using mini Quick Spin RNA Columns (Roche). RNAs were purified with phenol/chloroform and ethanol-precipitated. The RNA pellet was resuspended in RNAse-free water, and the final RNA concentration was quantified by measuring the absorbance at 260/280 nm. Binding reactions were carried out in 200 µL of SELEX buffer (20 mM MOPS pH 7.0, 50 mM KCl, 5LmM MgCl_2_, 1 mM DTT, 0.1% Triton-X-100, and 0.1LmM PMSF) and included 80 pmol of GST-fused protein immobilized on GSH beads (GE Healthcare), 1–8 µg of heparin sulfate, and 0.8–4 nmol of RNAs. Samples were gently agitated at 4L°C for 60 min. Unbound RNA was removed by five consecutive bead washings in 500 µL of SELEX buffer. HCl-25LmMLglycine (pH 2.2) was used to dissociate RNA-protein complexes (25 °C, 5 min). Dissociated RNAs were purified using mini Quick Spin RNA Columns (Roche), then ethanol-precipitated, and finally reverse-transcribed (SuperScript® III Reverse Transcriptase, Invitrogen). The resulting cDNAs were PCR amplified by 10 cycles for SELEX rounds 1 to 3, or 15 cycles for last SELEX round. Amplification products were then purified (PCR clean up kit, Thermo Fisher Scientific). The amplified cDNAs were used as a template to synthesize a new RNA library for the following SELEX rounds. In every SELEX round, selection stringency was intensified by increasing the RNA/protein ratios and the heparin concentration. The final PCR products (round 4) were subcloned into the pJET vector (CloneJET PCR Cloning Kit, Thermo Fisher Scientific) and sequenced. The obtained sequences were analyzed using the MEME tool (version 5.4.1) (Bailey *et al*., 2009), then logos were redesigned using the WebLogo software (Crooks *et al*., 2004).

### Structural modelling of multiprotein complexes involving SR45

Modelling of the 3D structure of different multiprotein complexes was performed with the AlphaFold software version 2.3.0 (Jumper *et al*., 2021). The multimer option was selected with a database version from January 1st, 2023, and a total of 25 models generated per prediction. All models were visually checked and the one with the best iptm+ptm (interface predicted template modelling + predicted template modelling) value was energy minimized within AlphaFold and selected for further analysis. Figures were generated with PyMOL v2.5.2 (The PyMOL Molecular Graphics System, Version 2.0 Schrödinger, LLC.).

### Statistical analysis

All data evaluation and statistics were performed using GraphPad Prism 7 (GraphPad Software v7.00). The replication level of each experiment is described in the Figure legends.

### Accession numbers and information

All gene sequences are available through *The Arabidopsis Information Resource* (TAIR, http://www.arabidopsis.org/), with the accession numbers: Arabidopsis *ACINUS* (AT4G39680)*, AFC1* (AT3G53570)*, AFC2* (AT4G24740)*, AFC3* (AT4G32660)*, ALY4* (AT5G37720)*, Cyp59* (AT1G53720)*, Cyp65* (AT5G67530)*, Cyp71* (AT3G44600)*, CypRS64* (AT3G63400)*, CypRS92* (AT4G32420)*, eIF4A3* (AT3G19760)*, GRP7* (AT2G21660)*, GRP8* (AT4G39260)*, MAGO* (AT1G02140)*, MOS12* (AT2G26430)*, MOS14* (AT5G62600)*, PININ* (AT1G15200)*, PRP38* (AT2G40650)*, RS2Z232* (AT3G53500)*, RS2Z33* (AT2G37340)*, RS31* (AT3G61860)*, RS31a* (AT2G46610)*, RS40* (AT4G25500)*, RS41* (AT5G52040)*, RSZ21* (AT1G23860)*, RSZ22* (AT4G31580)*, RSZ22a* (AT2G24590)*, RZ-1B* (AT1G60650)*, RZ-1C* (AT5G04280)*, SAP18* (AT2G45640)*, SC35* (AT5G64200)*, SCL28* (AT5G18810)*, SCL30* (AT3G55460)*, SCL30a* (AT3G13570)*, SCL33* (AT1G55310)*, SR30* (AT1G09140)*, SR34* (AT1G02840)*, SR34a* (AT3G49430)*, SR34b* (AT4G02430)*, SR45* (AT1G16610)*, SRPK1* (AT4G35500)*, SRPK2* (AT2G17530)*, SRPK3a* (AT5G22840)*, SRPK3b/SRPK5* (AT3G44850)*, SRPK4* (AT3G53030)*, Y14* (AT1G51510).

## Results

### SR45 is expressed at all developmental stages

During plant development, *SR45* was expressed to different extents in all vegetative and floral organs examined as described by quantitative RT-PCR profiling **(Figure 1A)**. *SR45* was significantly more expressed in inflorescences and young seedlings. The expression level was lower and similar in whole roots, stems and (cauline) leaves. We also determined the distribution of the protein using a GFP translational fusion (*pSR45:SR45-EGFP*) **(Figure 1B-R)** and the spatial expression pattern of *SR45* using a promoter:reporter construct (*pSR45:EGFP*) **(Supplementary Figure S3)**. Translational fusion lines confirmed (i) the spatial transcriptional activity of the *SR45* promoter and (ii) the nuclear localization of SR45 (Ali *et al*., 2003). The seed coat (including hilum) presented GFP fluorescence **(Figure 1B)**. Embryos freshly dissected from seeds displayed fluorescence in cotyledons **(Figure 1C)**. Right after germination, seedlings showed an intense fluorescence in all cell types within the radicle **(Figure 1D-E)**, the hypocotyl and the cotyledons **(Figure 1E)**. During vegetative growth, fluorescence was detected in roots, root hairs **(Figure 1E-G)**, and leaves (including trichomes and stomata) **(Figure 1I-J)** and was particularly visible within the vascular tissues of roots **(Figure 1F)**. During flowering, the receptacle at the flower bases, together with the abscission zone and the gynophore, emitted fluorescence signals **(Figure 1K)**. In flowers, GFP fluorescence resolved to styles and stigmas, stamens (anthers and filaments but not pollen grains), petals (including veins), and sepals **(Figure 1L-R)**. The observations in *pSR45:EGFP* lines were in agreement with those of the *pSR45:SR45-EGFP* translational fusion lines. However, fluorescence of GFP alone, which is not targeted to any specific subcellular compartment, was very weak, hence being sometimes hardly detected above the background **(Supplementary Figure S3)**. Indeed, seeds, embryos and radicles presented a low fluorescence signal in *pSR45:SR45-EGFP* lines that was fainter in *pSR45:EGFP* lines.

**Figure 1.**
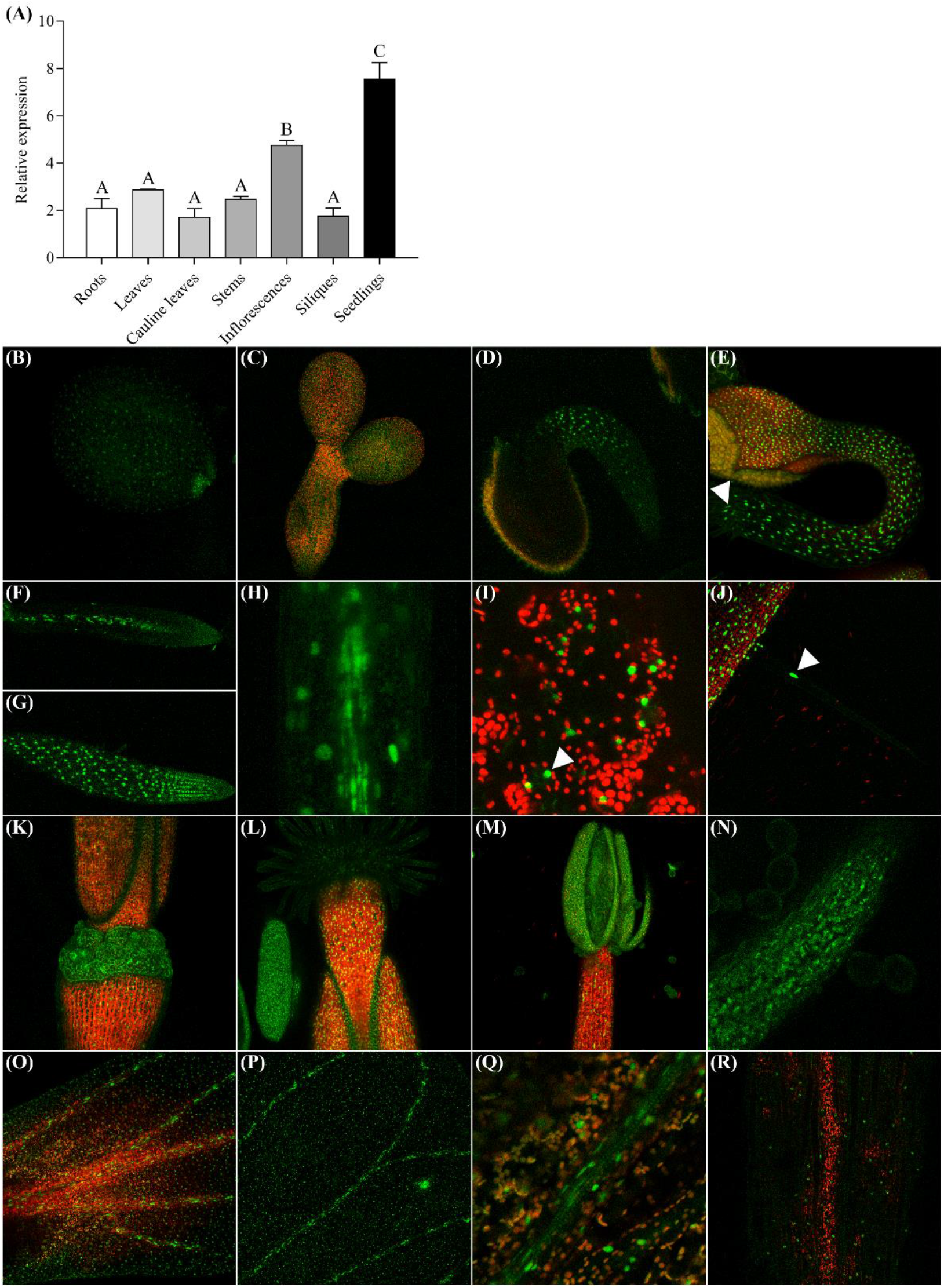
Expression profile of the *SR45* gene and of the corresponding protein. **(A)** Quantitative RT-PCR analyses of *SR45* gene expression in Arabidopsis vegetative and reproductive organs. Values represent means ± SEM (from three biological replicates, each consisting of pools of organs from three plants) and are relative to *AT1G58050*. Data were analyzed by one-way ANOVA followed by Dunnett multiple comparison post-tests. Statistically significant differences between means are indicated by different letters (P<0.05). (**B-R**) Expression profile of the SR45 protein. The fusion protein SR45:GFP is expressed in **(B)** seed coat and hilum region, **(C)** embryo (cotyledons), **(D)** radicle, **(E)** young seedling (cotyledons and root, including root hairs as indicated by the arrow), **(F-H)** two-week-old root vasculature **(F, H)** and epidermis **(G)**, **(I-J)** leaves, including stomata [**(I)**, arrow] and trichome [**(J)**, arrow], **(K-L)** gynoecium (valves, gynophore, abscission zone and receptacle) **(K)**, and stigma and style **(L)**, **(M-N)** androecium (anther and filament), **(O-Q)** petals (epidermis and veins), and **(R)** sepal epidermis. Red signals represent chlorophyll autofluoresence. At least three independent T3 lines were generated and analyzed, depicting similar fluorescence profiles.

### Structural determinants modulating SR45 localization

To understand the structural determinants modulating the SR45 cellular localization **(Figure 1)**, we engineered seven mutant variants **(Supplementary Figure S1)**. First, the canonical RRM of SR45, which is defined by a β_1_-α_1_-β_2_-β_3_-α_2_-β_4_ topology, was mutagenized at conserved residues mediating RNA recognition in both RNP1 and RNP2 motifs (Cléry *et al*., 2008). Both H101 (RNP2) and H141 (RNP1), together with Y143, were substituted to alanine to create the SR45mutRRM variant. Second, all Ser residues were substituted to Ala within the RS1 (nt 1 to 291, 39 S-to-A substitutions) and/or RS2 (nt 529 to 1245, 49 S-to-A substitutions), generating three variants (mutRS1, mutRS2, mutRS1+RS2), respectively. Finally, RRM and RS1/RS2 mutations were combined, producing three more mutant variants (mutRS1+RRM, mutRRM+RS2 and mutRS1+RRM+RS2).

After transient expression in tobacco leaf cells, no distinguishable changes appeared in the protein localization of the SR45mutRS1 or SR45mutRRM variants **(Figure 2)** compared to the native protein. In contrast, SR45mutRS2 partially accumulated in the cytoplasm in addition to a nuclear localization. These observations are in agreement with previous work examining the localization of RS1- or RS2-truncated SR45 mutants (Ali and Reddy, 2006; Ali *et al*., 2008). SR45mutRS1+RS2 showed nucleolar retention, whereas combining RS2 and RRM mutations (SR45mutRRM+RS2) noticeably increased the cytoplasmic retention compared to the native protein and the SR45mutRS2 variant. With all substitutions combined (SR45mutRS1+RRM+RS2), no fluorescence was detected within the nucleus, and only cytoplasmic signal was observed. Altogether, the integrity of the RS2 domain appears necessary but not sufficient for the nucleoplasmic localization of SR45.

**Figure 2.**
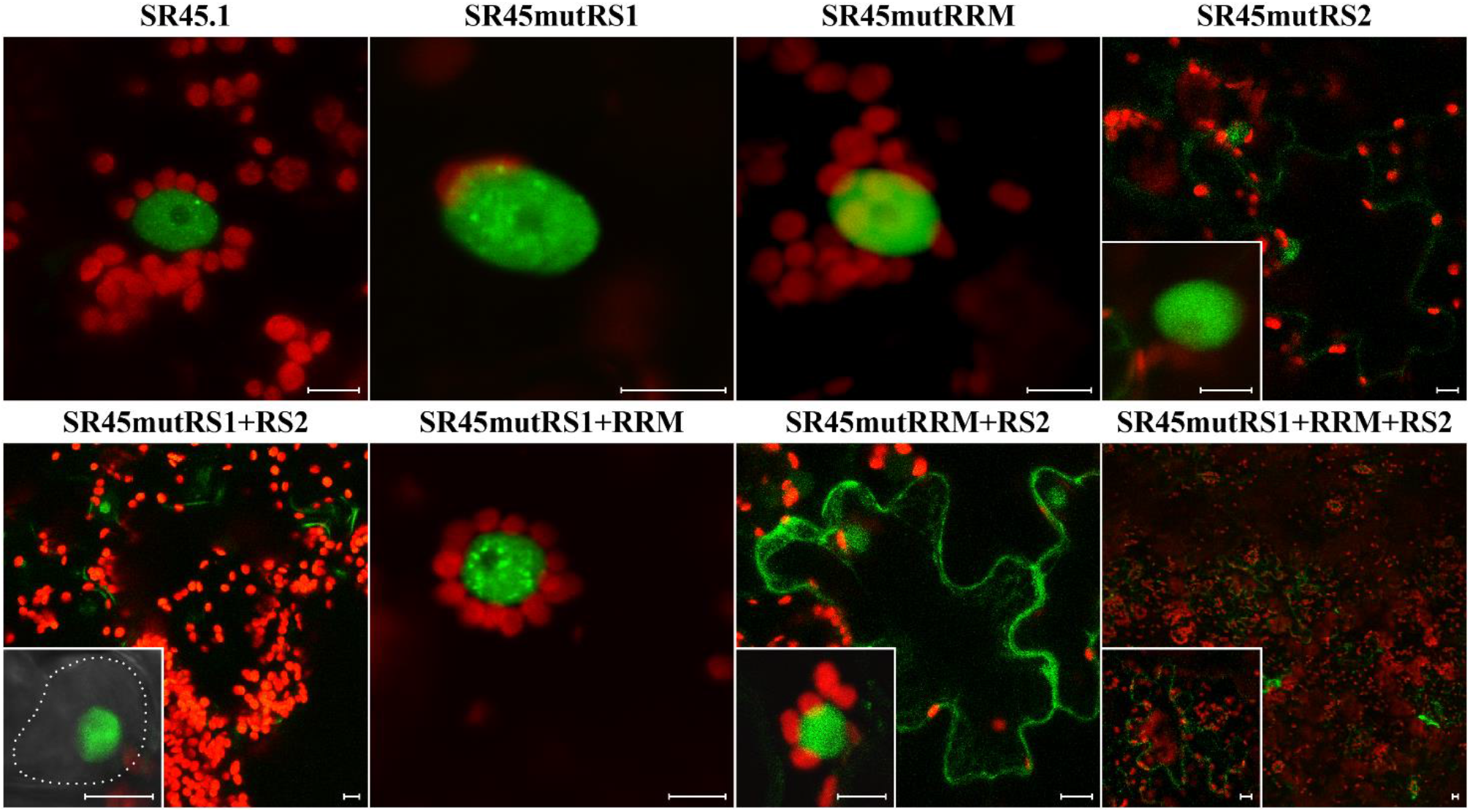
Structural determinants modulating SR45 localization. Subcellular fluorescence distribution in transient expression assays in tobacco leaf cells upon N-terminal EGFP-tagging of the native SR45 and SR45 mutant variants. Dotted lines delimit the nucleus. Scale bars = 10 µm. Red signals represent chlorophyll autofluoresence. Whenever cytoplasmic and nuclear signals were detected, insets depict the detailed intranuclear localization. At least three independent transient events were generated and analyzed, depicting similar fluorescence profiles.

### SR45 interactome

To identify interactors of SR45, two Arabidopsis cDNA libraries (a commercially available and a custom-made) were first screened using yeast two-hybrid (Y2H) and four previously unidentified interactors of SR45 were uncovered: CypRS64 (<u>Cy</u>clo<u>p</u>hilin-containing <u>RS</u> domain of <u>64</u> kDa), PRP38 (<u>P</u>re-m<u>R</u>NA <u>P</u>rocessing factor <u>38</u>), RS2Z32 (arginine/serine-rich zinc knuckle-containing protein 32), and MOS12 (*<u>M</u>ODIFIER <u>O</u>F <u>S</u>NC1-1*, <u>12</u>). These interactions were then further confirmed by directed yeast two-hybrid assays (dY2H) **(Figure 3**, **Supplementary Figure S4 and Supplemental Table S2)**.

**Figure 3.**
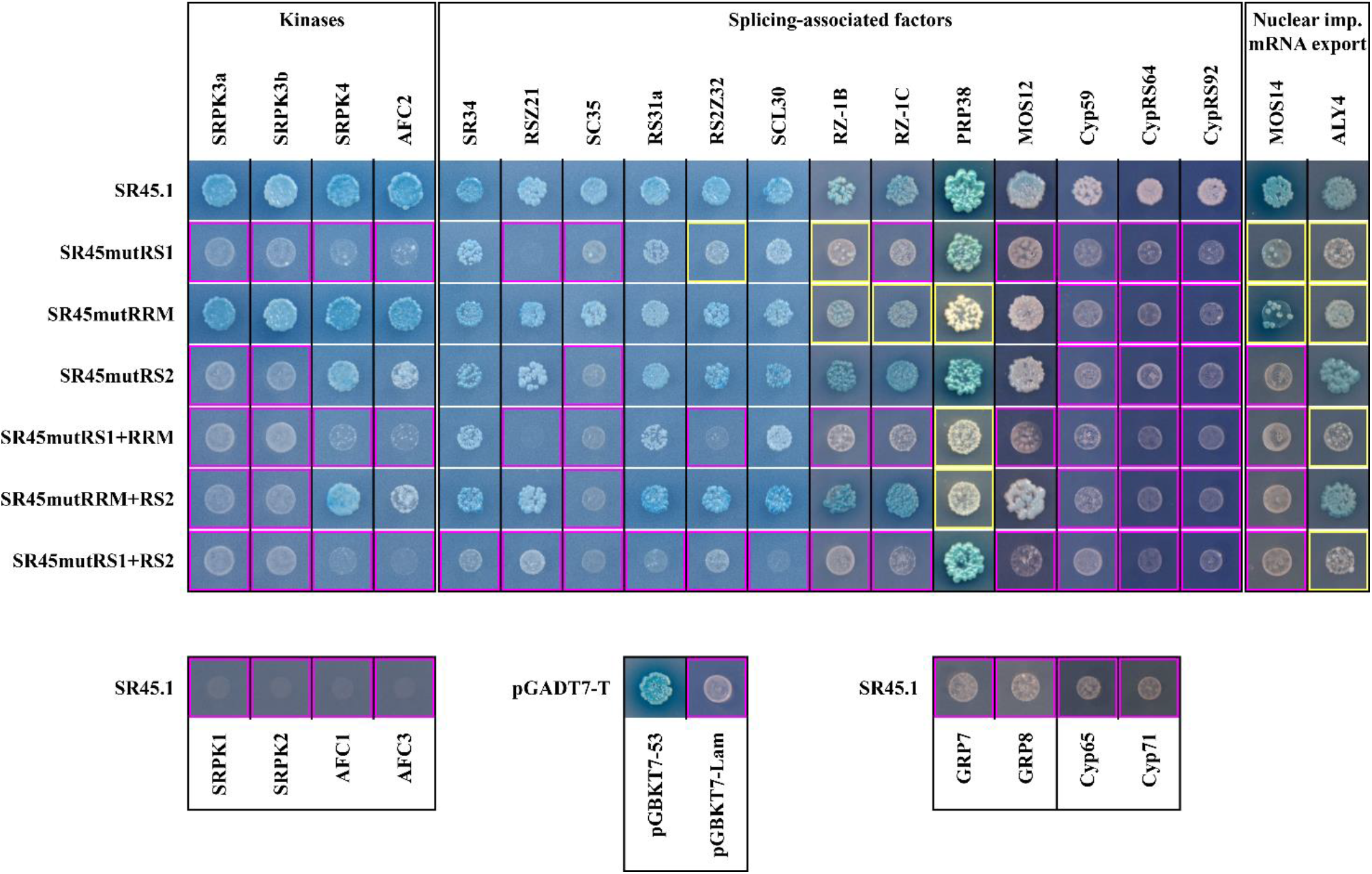
SR45 interactions detected by (directed) yeast two-hybrid screenings. From the mated culture, dilutions to an OD600 of 1 were spotted on -Trp/-Leu/-His/X-a-Gal/AurA agar plates. Positive interactions were confirmed by growth and blue staining as seen with the positive control (pGADT7-T x pGBKT7-53). Yellow and purple squares are indicative of a weakened or abolished interaction upon substitutions within the corresponding SR45 domain, respectively. A weakened interaction is characterized by the ability of several single colonies to grow, while an abolished interaction is characterized at most by a white halo of dead cells (similar to the negative control pGADT7-T x pGBKT7-Lam). An interpretive summary of the results is presented in **Supplementary Figure S4**.

dY2H assays were performed to examine the SR45 interactions with kinases potentially involved in RS domain phosphorylation, the 3 CLK kinases (AFC1-3) (Bender and Fink, 1994) and the 5 SRPK kinases encoded in the Arabidopsis genome (Lorković *et al*., 2004; de la Fuente van Bentem *et al*., 2006; Stankovic *et al*., 2016). We confirmed that SR45 interacts with AFC2 (Golovkin and Reddy, 1999), and confirmed its recently reported interactions with SRPK3a, SRPK3b [also named SRPK5 (Wang *et al*., 2023)] and SRPK4. Among all SR45 mutant variants, only RS1 mutations abolished the SRPK4- and AFC2-SR45 interactions, while RS1 or RS2 mutations both suppressed interactions with SRPK3a and SRPK3b/SRPK5 **(Figure 3)**. Those results suggested that RS domains are involved in interactions with kinases.

SR45 interacts with several other SR proteins, namely SCL33, RSZ21, SR30, SR34, SR34a, as well as other splicing factors such as the hnRNP-like proteins RZ-1B and RZ-1C (Golovkin and Reddy, 1999; Zhang *et al*., 2014; Stankovic *et al*., 2016; Wu *et al*., 2016). We showed here that SR45 was able to interact with all SR proteins but itself (SR45.1 or SR45.2 splicing isoforms (Zhang and Mount, 2009)) **(Supplementary Figure S6)**. The integrity of both RS domains was crucial for SR45 to contact SC35. In the cases of SR34, RS31a and SCL30, the interactions were only weakened by mutations applied to either RS1 or RS2 but were completely abolished for RS1/RS2 variants **(Figure 3)**. Instead, the RS1 variants showed suppressed interactions with RS2Z32, RSZ21, RZ-1B, RZ-1C and MOS12. In contrast to interactions of SR45 with SR proteins, the interactions with RZ-1B, RZ-1C or PRP38 were considerably weakened upon substitutions in the RRM domain **(Figure 3)**. The SR45-PRP38 interaction was, however, not affected by other substitutions **(Figure 3)**. SR45 contacted several nuclear cyclophilins including Cyp59, CypRS64 and CypRS92, but not Cyp65 and Cyp71, which are all involved in splicing regulation (Lorković *et al*., 2004; Gullerova *et al*., 2006; Barbosa Dos Santos and Park, 2019). However, none of the mutant variants was able to maintain interactions with any of the cyclophilins tested in yeast. No interaction was observed between SR45 and hnRNP-like proteins GRP7 and GRP8 **(Figure 3)**.

In animals, several shuttling SR proteins (SRSF1, SRSF3 and SRSF7) act as adaptors for mRNA nuclear export through the TAP/NXF1 (<u>N</u>uclear RNA e<u>x</u>port factor <u>1</u>) machinery and compete with ALY/REF for binding in the NXF1 complex (Huang *et al*., 2003). A gene encoding a homolog of the human TAP/NXF1 is missing in plant genomes (Merkle, 2011), but numerous ALY/REF orthologs were discovered in Arabidopsis (Pfaff *et al*., 2018; Ehrnsberger *et al*., 2019). Our dY2H analysis highlighted an interaction between SR45 and ALY4, which was tremendously weakened by RS1 and/or RRM mutations **(Figure 3)**. SR45 presents a nucleocytoplasmic dynamic whose nuclear export is at least partly controlled by the XPO1-dependent export pathway (Stankovic *et al*., 2016), whereas the nuclear import of plant SR proteins is likely driven by the activity of MOS14 (*<u>M</u>ODIFIER <u>O</u>F <u>S</u>NC1-1*, <u>14</u>), a transportin-SR (TRN-SR) (Xu *et al*., 2011). Here, SR45 and MOS14 interacted, and the SR45 RS2 domain was required **(Figure 3)**.

The human RNPS1 contributes, together with SAP18, ACINUS and PININ, to the formation of two EJC peripheral complexes, ASAP and PSAP (Sakashita *et al*., 2004; Singh *et al*., 2010; Murachelli *et al*., 2012). We next performed dY2H and directed yeast three-hybrid (dY3H) assays to investigate interactions between SR45 with core (Y14, MAGO and eIF4AIII) and peripheral (SAP18 and either ACINUS or PININ, which are part of the ASAP and PSAP complexes, respectively) components of the EJC **(Figure 4 and Supplementary Figure S5)**. We demonstrated that, unlike eIF4AIII, MAGO and Y14 were able to interact with SR45, which required an intact RS1 domain **(Figure 4A)**. We further confirmed that SR45 was able to interact with both ACINUS and PININ (Qüesta *et al*., 2016; Chen *et al*., 2019; Bi *et al*., 2021) but not SAP18 (Wang *et al*., 2023). SAP18 was not able to interact with either ACINUS or PININ **(Figure 4A)**, however, when the three components were co-expressed in a Y3H system, an interaction was established between SR45 and SAP18 through either ACINUS or PININ **(Figure 4B)**. The SR45-ACINUS interaction was greatly diminished upon mutations in the RS1 domain **(Figure 4A)**. Based on animal models where two conserved residues, a threonine and a tyrosine, are critical for the assembly of the ASAP complex (Sakashita *et al*., 2004; Singh *et al*., 2010; Murachelli *et al*., 2012), we identified two amino acids (Thr607 and Tyr615) within the C-terminal extension of the RRM domain of ACINUS that were expected to be critical for the interaction with SR45 **(Supplementary Figure S7D)**. Accordingly, mutations of these two residues (T607A/Y615A) abolished the interaction of ACINUS with SR45. On the other hand, the SR45-PININ interaction was suppressed when mutations were present in the RRM of SR45 **(Figure 4A)**.

**Figure 4.**
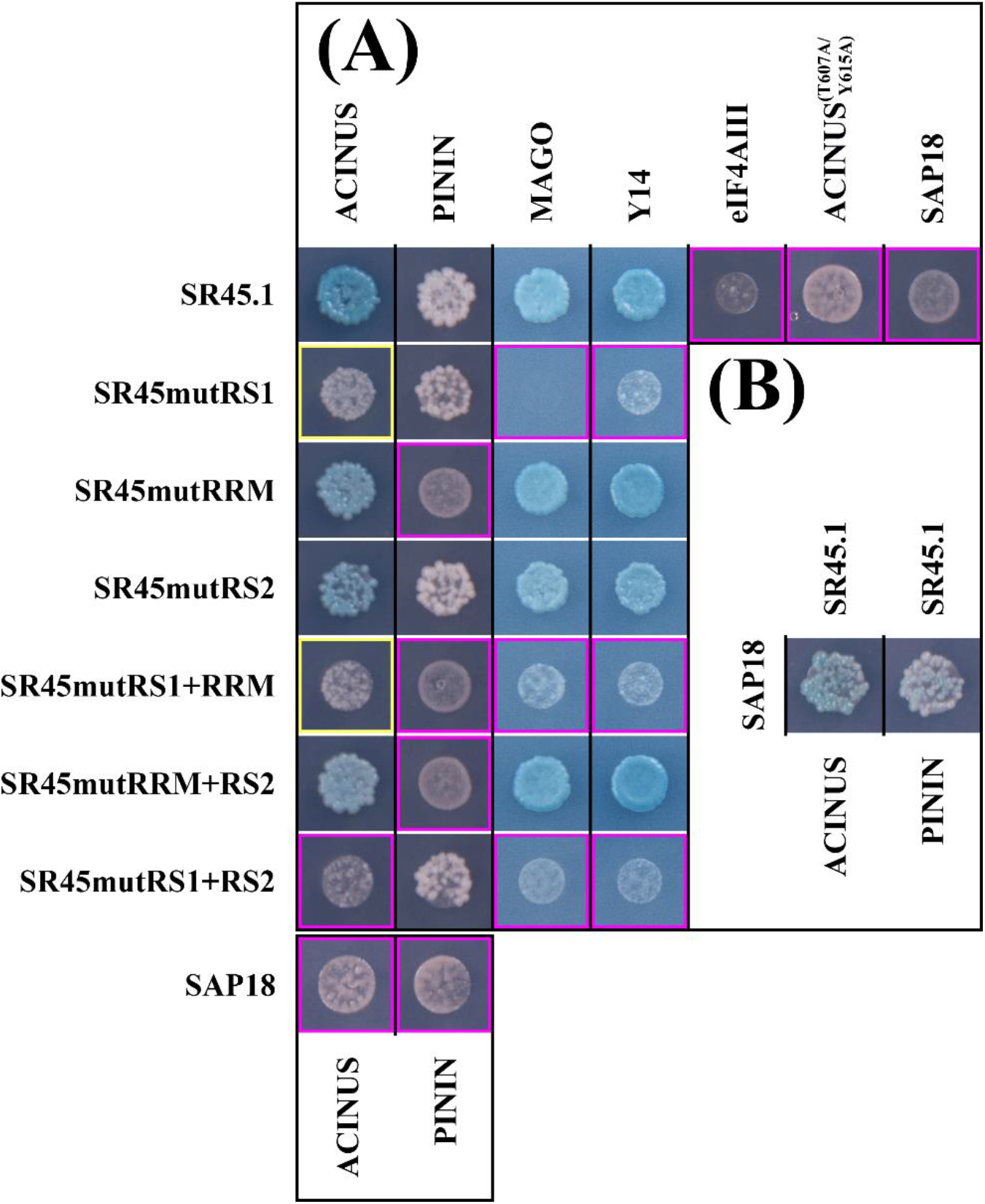
SR45 associates with EJC core components and peripheral factors. Interaction between the native SR45, and SR45 mutant variants, and ACINUS(RRM) and PININ (from plant ASAP and PSAP complexes, respectively), and EJC core components (MAGO and Y14) **(A)**. Ternary complexes formed in the simultaneous presence of SAP18, SR45 and either ACINUS(RRM) or PININ **(B)**. From the mated culture, dilutions to an OD600 of 1 were spotted on -Trp/-Leu/-His/X-a-Gal/AurA agar plates (for Y2H assays) **(A)** or -Met/-Trp/-Leu/-His/X-a-gal/Aureobasidin A (for Y3H assays) **(B)**. Positive interactions were confirmed by growth and blue staining. Yellow and purple squares are indicative of a weakened or abolished interaction upon substitutions within the corresponding SR45 domain, respectively. A weakened interaction is characterized by the ability of several single colonies to grow, while an abolished interaction is characterized at most by a white halo of dead cells. An interpretive summary of the results is presented in **Supplementary Figure S5**.

Altogether, our observations confirmed the importance of serine residues in the establishment of interactions with kinases and splicing factors but also depicted a role for the canonical RRM (through both RNP1 and RNP2 motifs) in the formation of a protein complex involved in mRNA export and mRNA surveillance.

### Structural modelling of multiprotein complexes involving SR45

Structural modelling was conducted to further explore the details of the interactions taking place in two multiprotein complexes involving SR45: ACINUS-SR45-SAP18 (ASAP) and PININ-SR45-SAP18 (PSAP) **(Supplementary Data S1 and S2)**.

For ASAP, clear interactions were observed between structured regions of the three partners, which all contained extended unstructured regions with predicted local distance difference test (pLDDT) values below 50 (ACINUS residues 1-3, 66-444 and 542-597; SR45 residues 1-95 and 177-414; SAP18 residues 1-28). For SR45, this confirmed the unstructured nature of the RS1 (1-97) and RS2 (177-414) domains, surrounding a structured RRM domain (98-176) **(Supplementary Figure S1)**. Outside of these unstructured regions, AlphaFold’s estimation of the accuracy was high with an average pLDDT of 89.5 **(Supplementary Figure S7A-B)**. The ACINUS N-terminal SAP domain (residues 4-66) was well folded but did not interact with any other part of the complex, and will not be further discussed. The interactions between the RRM domain of SR45, the ubiquitin-like domain of SAP18 and the last 35 residues of ACINUS were very similar to that observed in the ASAP complex available in the Protein Data Bank (PDB code 4A8X) (**Supplementary Figure S7B-C**) with a Root-Mean-Square Deviation (RMSD) of 1.0Å calculated over 201 Cα of the 218 residues spanning the three chains of the complex (Murachelli *et al*., 2012). The sequence identity between residues present in this structure (human RNPS1, ACINUS peptide from *Drosophila melanogaster* and *Mus musculus* SAP18) and equivalent residues in the Arabidopsis model was 49%. The model of the core ASAP structure confirmed that (i) ACINUS interacted with SR45 through two residues (T607 and Y615) located in the vicinity of its RRM, and that (ii) the three substituted residues in the RRM of SR45 were not involved in this interaction, corroborating Y2H assays **(Figure 4A)**. In addition to this known part of the ASAP complex in animals, the AlphaFold model revealed a potential additional interaction between the RRM domain of ACINUS (residues 445 to 541) and the ubiquitin-like domain of SAP18. The interface, which covers an area of 588Å^2^, involved residues from the L1, L3 and L5 loops **(Supplementary Figure 7D)**, the first two being the most conserved regions on the surface of the domain according to Fernandes and colleagues (Fernandes *et al*., 2018). With only two salt bridges and some hydrophobic interactions, the interaction between the two partners was not strong according to the PISA server (Krissinel and Henrick, 2007) and could potentially be strengthened upon interactions with other partners including RNA.

In the PSAP complex, which is characterized by an average pLDDT value calculated for structured region (pLDDT > 50) of 86.5, the same interactions are present between the ubiquitin-like domain of SAP18 and the RRM domain of SR45, and as predicted by Murachelli and colleagues (Murachelli *et al*., 2012), the ^310^KAEPRIYYAPVKPL^323^ peptide of PININ mostly substituted the C-terminal domain of ACINUS at the junction between SAP18 and SR45 in the ASAP complex **(Supplementary Figure 8A-C)**. In addition to this peptide, the structured part of this model of PININ was made of four α helices (α1-α4) with pLDDT value over 50. Three of these helices were approximately parallel: α1 (residues 8-32), α3 (residues 188-258) and α4 (residues 327-393). The α2 helix (residues 160-186) directly preceded α3 and was perpendicular to the three other helices, but the confidence of the model was relatively weak in this region. The α3 and α4 helices, which were separated by a loop containing the ACINUS-like peptide and a 50 amino acids long unstructured stretch, also interacted with the ubiquitin-like domain of SAP18 **(Supplementary Figure 8D)** and contributed to the interaction with three H-bonds, two salt bridges and few hydrophobic interactions, covering a 764Å^2^ surface area. The four α helices might provide an extended platform for the recruitment of multiple additional partners.

### The RS1 and RS2 domains of SR45 connect splicing factors and EJC peripheral factors in planta

Observed in yeast and in the ASAP structural model described above, the interactions underlying the involvement of SR45 into the splicing reaction (ACINUS, PININ and SR34) as well as the spliceosome catalytic activation (PRP38) (Schütze *et al*., 2016), which are also established by RNPS1 in human (Schütze *et al*., 2016; Schlautmann *et al*., 2022), were further characterized in living plant cells using Yellow Fluorescent Protein (YFP)-based bimolecular fluorescence complementation (BiFC). BiFC experiments were performed in tobacco leaf cells where the native or mutant variants of SR45 fused to the C-terminal fragment of YFP (^C^YFP) and interactors fused to the N-terminal fragment of YFP (^N^YFP) were transiently co-expressed. YFP fluorescence was observed in the nucleus and in structures similar to nuclear speckles, suggesting that SR45 interactions with ACINUS, SR34 and PRP38 occurred in the nucleus and in splicing factors-enriched granules *in planta* **(Figure 5A)**. The study of mutant variants confirmed results obtained in yeast cells **(Figure 3)**, as SR45mutRS1+RS2 was unable to interact with ACINUS and SR34, but remained able to establish an interaction with PRP38 within the nucleolus. Furthermore, mutations in the RS1 or RRM domains did not affect any of the SR45 associations mentioned above, similarly to what was observed in yeast cells and in the modelling performed for ACINUS **(Supplementary Figure S7)**. Trimolecular fluorescence complementation (TriFC) also confirmed dY3H assays as SAP18 was unable to interact with either SR45, ACINUS or PININ **(Figure 5B, upper panels)** but was able to form trimeric ASAP and PSAP complexes **(Figure 5B, lower panels)**.

**Figure 5.**
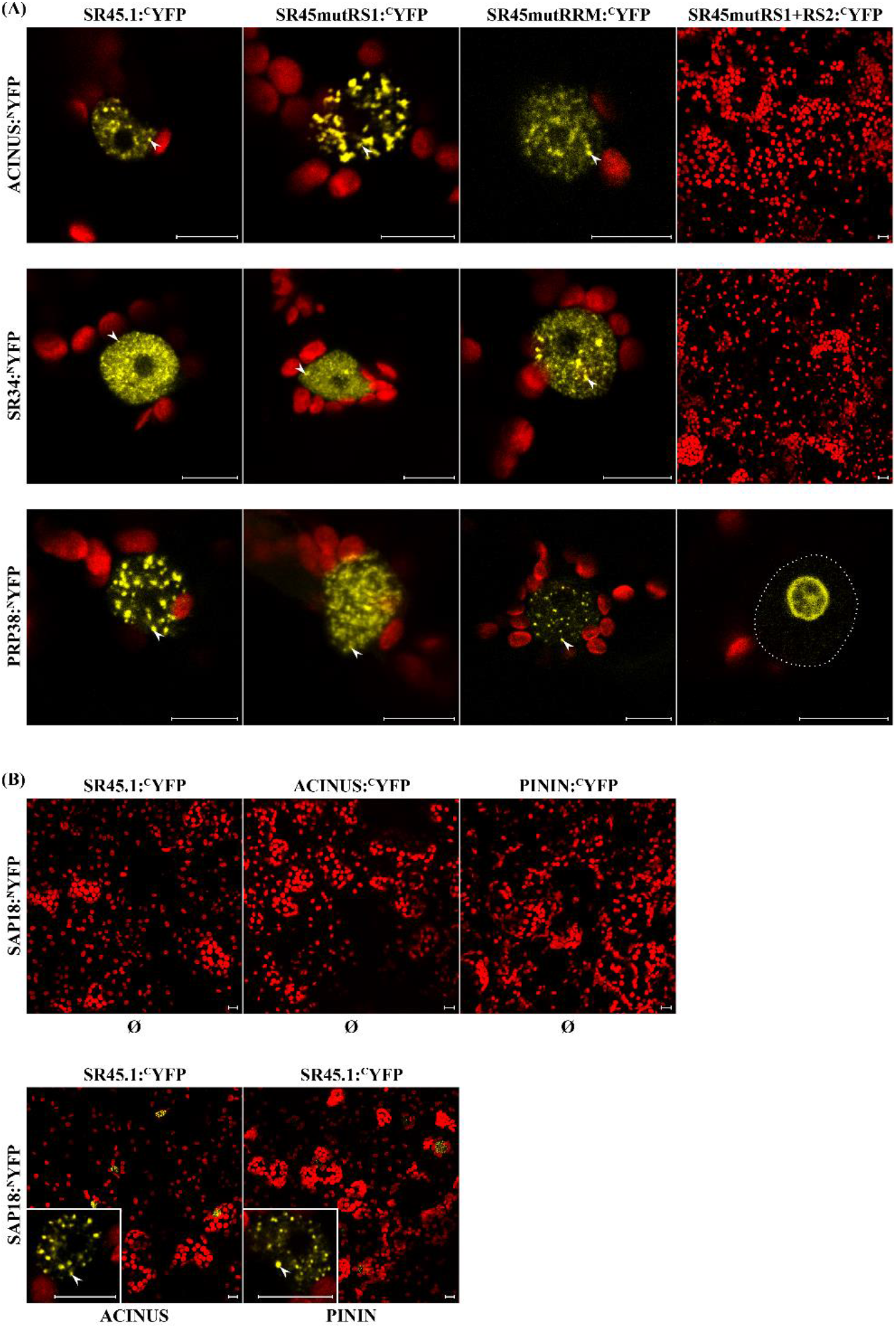
SR45 interacts with splicing factors and EJC peripheral factors in plant cells. **(A)** Bimolecular Fluorescence Complementation (BiFC) through co-expression of SR45 or SR45 mutant variants fused to the C-terminal half of YFP (SR45, SR45mutRS1, or SR45mutRS1+RS2:^C^YFP) and ACINUS(RRM), SR34, or PRP38 fused to the N-terminal half of YFP (ACINUS(RRM), SR34, or PRP38:^N^YFP). Dotted lines delimit the nucleus. **(B)** Trimolecular Fluorescence Complementation (TriFC) through co-expression of SR45:^C^YFP, ACINUS:^C^YFP or PININ:^C^YFP with SAP18:^N^YFP and a pMDC32 plasmid harboring no coding sequence (Ø, upper panels) or through co-expression of SR45:^C^YFP with SAP18:^N^YFP and a pMDC32 harboring the coding sequence of either the RRM of ACINUS (bottom left panel) or PININ (bottom right panel). Pictures represent the maximum intensity projections of 30 z-stack images from a 29.21 μm section (with a z-step size of 1.01 µm). The voltage of the EYFP PMT was set up at 700V in all experiments. For both BiFC **(A)** and TriFC **(B)** observations, representative images of fluorescence reveal the interaction of SR45 variants in the nucleus, nucleolus, and in speckles-like structures (arrowheads) in transient expression assay in tobacco leaf cells. At least three independent transient events were generated and analyzed, depicting similar fluorescence profiles. Scale bars = 10 µm. Red signals represent chlorophyll autofluoresence.

### Subcellular localization of SR45-associated partners

To better decipher the dynamic phosphorylation of SR45 across cellular compartments and where SR45 might precisely contact its (peripheral) EJC interactors in plant cells, EGFP was fused to the N-terminus of candidate proteins, and translational fusions were transiently expressed in tobacco leaf cells. SRPK1 and SRPK2, two non-interactors of SR45, resided exclusively in the cytoplasm **(Figure 6)**. Unlike SRPK4 which resided predominantly in the cytoplasm (alongside a weak detectable signal within the nucleus), SRPK3a and SRPK3b/SRPK5 localized in the cytoplasm and the nucleus **(Figure 6)**. The CLKs were the only kinases studied that accumulated in nuclear speckles-like structures **(Figure 6)**. ACINUS, SAP18 and PININ all localized in the nucleus, however, SAP18 also displayed a cytoplasmic localization **(Figure 6)**. Similarly, MAGO and Y14 were also detected in the nucleus and in the cytoplasm **(Figure 6)**. PRP38 localized in the nucleoplasm, speckles-like structures and the nucleolus **(Figure 6)**.

**Figure 6.**
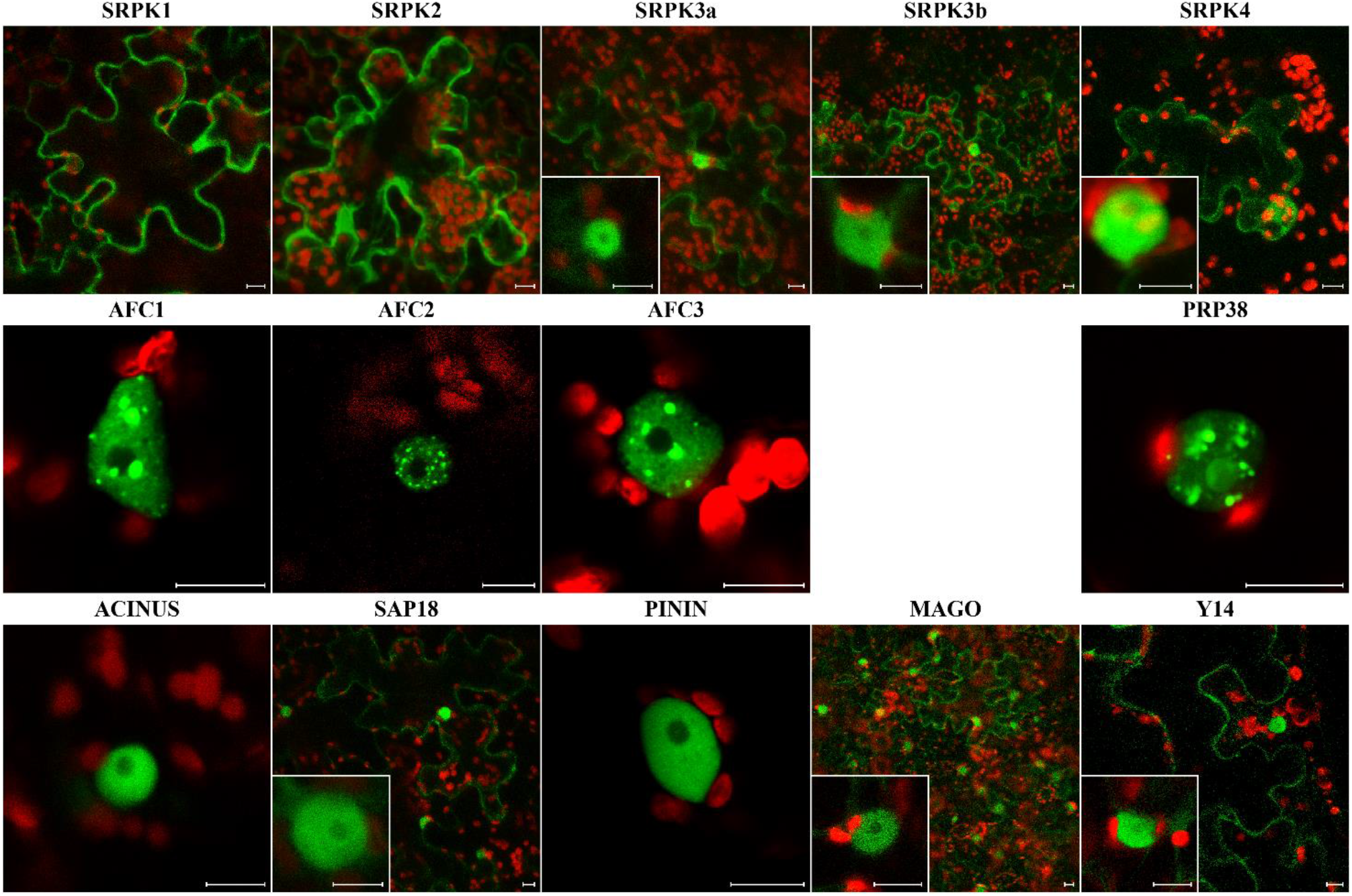
Subcellular localization of SR45-associated proteins. **(A)** Subcellular fluorescence distribution in transient expression assays in tobacco leaf cells upon N-terminal EGFP-tagging (except for SRPK4 which was C-terminally tagged). Scale bars = 10 µm. Red signals represent chlorophyll autofluoresence. Whenever cytoplasmic and nuclear signals were detected, insets depict the detailed intranuclear localization. At least three independent transient events were generated and analyzed, depicting similar fluorescence profiles.

These observations suggest that the interactions described by dY2H/dY3H and BiFC/TriFC may actually take place in plant cells, as the intracellular localization of SR45 and its kinase and EJC interactors at least partially overlapped.

### SR45 is binding to purine-rich RNA motifs

Then, we investigated the RNA-binding specificity of a recombinant RRM domain of SR45 fused to a glutathione-S-transferase using Systematic Evolution of Ligands by EXponential enrichment (SELEX). Among the twelve clones sequenced after SELEX, two statistically significant consensus sequences recognized by the SR45 RMM were identified, with a specificity for a purine-rich central motif (5’-[A/G]GAAG-3’) flanked by pyrimidine-rich regions (5’ UC[C/U] and 3’ UU[C/U][C/U]C) (*E*-value: 8.6 x 10^-4^) **(Figure 7A)** or for a purine-rich motif ([A/G]GA[C/A]A) (*E*-value: 1.2 x 10^-10^) **(Figure 7B)**. Because of the higher variability of bases at most positions in the 2^nd^ motif, we analyzed the 4-mer and 5-mer frequencies among the sequences used to identify it. The analysis of 4-mer frequency showed a higher occurrence of the GACA and GAAG motifs. The analysis of 5-mer frequency demonstrated higher prevalence of AAGUU, AGACA, GAAGU, GACAU and AGAAG motifs **(Supplementary Table S10)**. This is strikingly similar to G/A-rich motifs bound by RNPS1 (Mabin *et al*., 2018).

**Figure 7.**
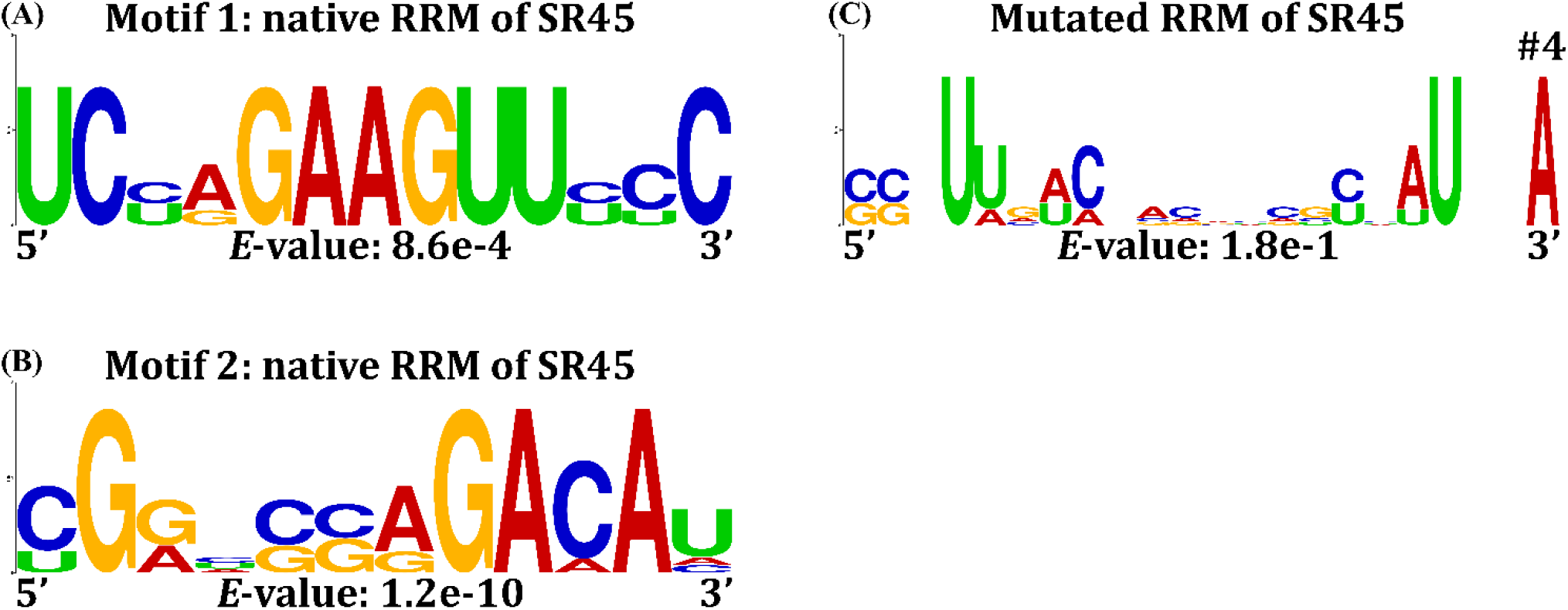
RNA motifs binding to the SR45 RRM. RNA motif 1 **(A)** and RNA motif 2 **(B)** identified through 4 rounds of SELEX selection with the RRM of SR45. Non-significant RNA motifs **(C)** identified using the mutated version of the RRM of SR45. The statistical significance (*E*-value) is indicated at the bottom of each consensus. Logos were redesigned using WebLogo (Crooks *et al*., 2004). The fourth non-significant RNA motif (#4) identified using the mutated version of SR45 RRM (which presents the best *E*-value) is displayed here, but the three other RNA motifs (#1-3) are depicted in **Supplementary Figure S9**. All RNA motifs were discovered using the MEME tool (version 5.4.1).

To ensure the specificity of the binding of the identified RNA motifs to the SR45 RRM, a mutated version (H101A, H141A and Y143A) was produced and studied in the SELEX set-up. Four different sets of twelve sequences obtained using the mutated SR45 RRM were analyzed **(Supplementary Table S11)** using the MEME tool. No enriched motif was discovered **(Figure 7C and Supplementary Figure S9)**, and the best motif recovered (lowest *E*-value, #4) did not resemble significant motifs found for the native canonical RRM **(Figures 7A-B)**, indicating that amino acids (H101, H141 and Y143) mutated within the RRM are necessary and sufficient to ensure the accurate binding of SR45 to its purine-enriched target motif.

## Discussion

Pre-mRNA splicing is a crucial step in the regulation of gene expression. Splice-site selection is carried out by the binding of many *trans*-acting protein factors including the SR splicing factors (Chen and Manley, 2009). In mammalian cells, it has been shown that SR proteins regulate many aspects of gene expression (Huang and Steitz, 2001). In contrast, their plant counterparts are mainly known to play important roles in constitutive splicing and alternative splicing, in biological processes such as ABA signalling, salt response, metal homeostasis, or response to high-light (Xing *et al*., 2015; Yan *et al*., 2017; Zhang *et al*., 2017; Fanara *et al*., 2022; Albaqami, 2023) although they are also suspected to be involved in other processes regulating gene expression, e.g. RNA-directed DNA methylation and regulation of transcription (Ausin *et al*., 2012; Qüesta *et al*., 2016; Yan *et al*., 2017; Bi *et al*., 2021; Mikulski *et al*., 2022).

### The RS domains of SR45 control its subcellular localization and protein-protein interaction network

The SR45 splicing factor displays a modular structure, with two unstructured RS and one structured RRM domains, which is atypical for a plant SR protein (Barta *et al*., 2010; Califice *et al*., 2012). In this report, we examined the involvement of each of these domains, as well as some of their constituent amino acid residues, in molecular functions known to be regulated by the human ortholog of SR45 (RNPS1) (Califice *et al*., 2012), whose association to EJC peripheral factors determines its roles in pre-mRNA splicing, mRNA export and quality control (Le Hir *et al*., 2016) **(Figure 8)**. To this avail, all serine residues were substituted into alanines in the RS domains and three point mutations were introduced in conserved residues of the RRM domain. One might question the strategy of modifying multiple amino acids and its impact on the global structure of SR45, hence on the functionality of the engineered mutant variants. We studied here the impact of these substitutions on (i) the protein localization, demonstrating that all mutant variants displayed a complete or partial nuclear localization similarly to the native protein (with the exception of the SR45mutRS1+RRM+RS2 that was not further studied); (ii) the protein-protein interactions, for which we demonstrated that the mutations had different impacts on the interactions with different partners in yeast cells and *in planta*, dismissing the possibility of a complete non-functionality of the mutant variants, and (iii) the protein-RNA interactions, demonstrating that the mutated RRM was still able to engage in interactions with RNA molecules despite an alteration of specificity, as it is well documented (Yang *et al*., 2011; Pabis *et al*., 2019; Ni *et al*., 2023).

**Figure 8.**
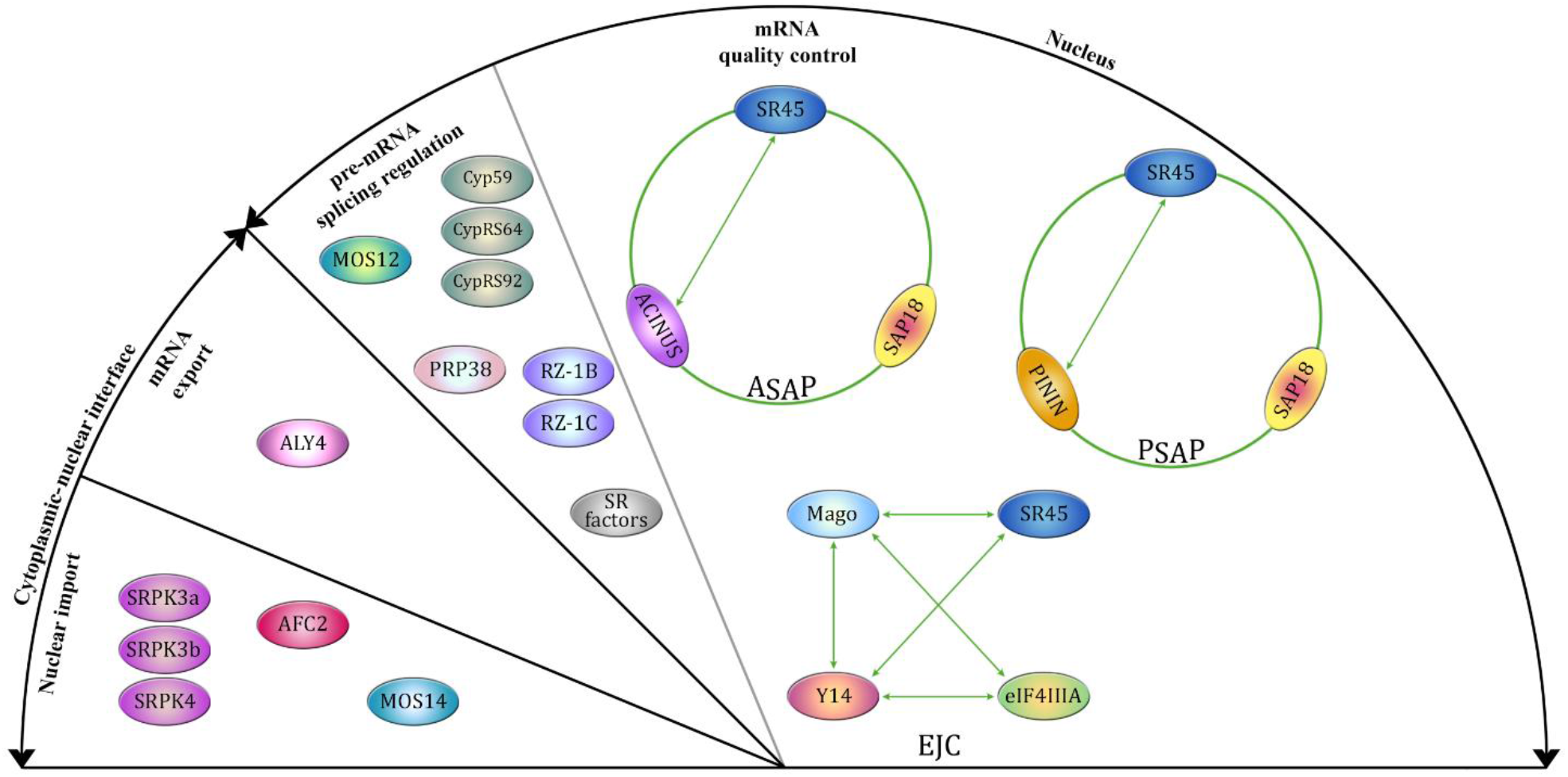
A working model for the molecular function of SR45-binding proteins. Schematic diagram showing the different binding partners of SR45 identified in this study and the functional implications of these interactions. Green and red double-headed arrows indicate, respectively, the presence or the absence of interaction during directed Y2H experiments. Green circles represent positive trimeric complexes observed by Y3H experiments and TriFC *in planta*. The grey divider represents functional interconnection between nuclear complexes. The absence of interaction discussed in this study is not displayed [e.g., between either SR45, ACINUS or PININ and SAP18, or between SR45 and either eIF4IIIA and ACINUS(T607A/Y615A)].

With this experimental set-up, we show that a number of interactions between SR45 and spliceosomal or EJC components entirely rely on the presence of serine residues in the RS domains of SR45. Indeed, the substitution of only few residues in these domains can either tremendously diminish or abolish the connection between SR45 and its partners, for instance the interactions with SR34 and other SR proteins. Some of those serine residues also control the subcellular localization of SR45. S-to-A substitutions within the RS2 domain alone induce cytoplasmic accumulation of SR45, which might result from its inability to contact the plant transportin-SR (MOS14) that is thought to be involved in the nuclear import of plant SR proteins (Xu *et al*., 2011) as its human ortholog TNPO3 (Maertens *et al*., 2014; George *et al*., 2019; Costa *et al*., 2021). In contrast, S-to-A substitutions in both RS domains strictly restrict SR45 to the nucleolus where it still can interact with PRP38. The subcellular localizations of RS mutant variants reported here refine the observations made by Ali and colleagues (2006 and 2008), revealing that the accumulation of SR45 within the nucleus or in both the nucleus and cytoplasm upon, respectively, disruption of the RS1 (SR45mutRS1 in this study) or RS2 domains (SR45mutRS2 in this study) are dependent of serine residues (Ali and Reddy, 2006; Ali *et al*., 2008). The nuclear import of SR proteins further depends on the phosphorylation of serine residues present in the RS domain (Lai *et al*., 2000; Lai *et al*., 2001). This process involves several kinases in animals, mainly SRPK1 (ortholog of the Arabidopsis SRPK4) and CLK1 (ortholog of the Arabidopsis AFC2), which act together to regulate the phosphorylation status of the human SRSF1 SR protein (Aubol *et al*., 2016). We show here that in Arabidopsis, the recognition of SR45 by two kinases of the SRPK family (SRPK3a and SRPK3b/SRPK5) depends on the integrity of the RS2 domain. These kinases might therefore be predominately involved (over other kinases) in the nuclear import of SR45 in conjunction with the transportin-SR MOS14. While S-to-A substitutions within the RS2 does not affect the association between SR45 and SRPK4 or AFC2, all mutant variants harbouring RS1 substitutions were not able to contact any of the kinases identified in this study. It is thus tempting to speculate that the RS1 domain of SR45, which is essential for AFC2 and SRPK4 binding, might serve as a platform for coordinating activity of these two kinases as seen in SRSF1 (Aubol *et al*., 2017; Wegener and Müller-McNicoll, 2019). A recent study identified specific serine residues within both RS domains whose phosphorylation levels were downregulated in a *srpk3 srpk4 srpk3b/srpk5* triple mutant, and tested interactions between these three kinases and SR45 mutant variants harbouring phosphoresidue substitutions to alanine (Wang *et al*., 2023). Here we focused on substituting all serines within a domain to better decipher the importance of the global phosphorylation status to (i) the subcellular localization of SR45 and (ii) to the protein-protein interaction network established by this splicing factor. Both approaches uncovered similar results, as SRPK3, SRPK4 and SRPK3b/SRPK5 interactions with SR45 involve native serine residues within both RS domains, i.e. S-to-A substitutions of all serines (this study) or of only phosphorylated serines (Wang *et al*., 2023) lead to the disruption of these interactions. The triple mutation *srpkii* notably reduces the phosphorylation level of SR45 at the T264 residue (Wang *et al*., 2023). This specific residue was in addition shown to be involved in the ABA-controlled phosphodegradation of SR45 (Albuquerque-Martins *et al*., 2023), suggesting a connection between the SRPK proteins, the stability of their SR45 substrate and ABA signalling.

### The SR45 RRM is essential to contact protein partners

In the RRM of SR45, a histidine residue is present at the position of the putative aromatic residue contacting RNA in both RNP2 (2^nd^ position) and RNP1 (3^rd^ position) motifs (Maris *et al*., 2005; Cléry *et al*., 2008; Daubner *et al*., 2013) **(Supplementary Figures S1 and S7D)**. Because histidine residues are known to bind RNA molecules (Morozova *et al*., 2006; Goroncy *et al*., 2010), both H101 (RNP2) and H141 (RNP1) together with Y143 were substituted by alanine to engineer the SR45mutRRM variant. The data presented here suggest that the H101, H141 and Y143 residues in the SR45 RRM are not structural determinants modulating SR45 localization but they are partially involved in SR45 association to RZ-1B, RZ-1C, ALY4 and PRP38. According to the structural models obtained in this study **(Supplementary Figures S7 and S8, and Supplementary Data S1 and S2)**, however, these residues are not involved in interactions with the two other partners in the ASAP and PSAP complexes. While the suppression of contact between the mutated RRM domain of SR45 and PININ is not supported by the model obtained with AlphaFold, the structural models are in agreement with a previous report establishing the involvement of the RRM of RNPS1 in the interaction with ACINUS and PININ (Murachelli *et al*., 2012). Although RNPS1 forms the ASAP complex with ACINUS and SAP18 through its RRM (Murachelli *et al*., 2012), SR45 contacts ACINUS using preferentially the RS1 domain. This interaction between unstructured regions, which is not detected in the model, is completely abolished when S-to-A substitutions are applied to both RS domains and is in agreement with recent work (Wang *et al*., 2023). The model of the core ASAP structure confirms that ACINUS contacts SR45 through two residues (T607 and Y615) located in the C-terminal extension of its RRM, supporting our Y2H data **(Figure 4A)**. Accordingly, the substitutions T607A and Y615A abolished the interaction between both partners in yeast. Our observations indicate that in addition to the core ASAP structure involving the SR45 RRM, the SAP18 ubiquitin-like domain and the ACINUS C-terminal peptide known in animals (i) the ASAP complex involves the RRM of ACINUS according to the model and at least the RS1 domain of SR45 according to Y2H observations, and (ii) the ASAP complex involves SAP18 as a ternary partner of a pre-existing SR45-ACINUS platform as shown by Y3H and TriFC assays.

### RNA binding specificity of the SR45 RRM

So far, the characterization of RNA-binding specificity of plant SR proteins mainly involved *in silico* analyses performed on RNA-Seq data generated from loss-of-function mutants. Such an approach suggested that SCL30, one of the interactors of SR45, was able to bind to a specific RNA sequence, 5’-AGAAGA-3’ (Yan *et al*., 2017). Four overrepresented RNA motifs were also previously identified by the identification of direct RNA targets of SR45 (SR45-associated RNAs, SARs) in seedlings (Xing *et al*., 2015): the identified M1 and M4 motifs are G/A-rich and were predicted to be predominately positioned within exons (Xing *et al*., 2015); the M2 and M3 motifs were U/C-rich and were preferentially located in intronic regions (Xing *et al*., 2015). While the “Motif 1” identified here through SELEX cycles combined features of all the motifs previously identified for SR45 (Xing *et al*., 2015), the “Motif 2” contained invariant G/A nucleotides and tend to be enriched in AGA, GGA and UCC trinucleotides **(Supplementary Figure S10)**. The human RNPS1 was found to be associated to G/A-rich 6-mers RNA regions (Mabin *et al*., 2018), which is reminiscent of the invariant sequence 5’-[A/G]GAAG-3’ found here in “Motif 1” associated to SR45. The RNA-binding specificity of the splicing factor and protein partner of SR45, RZ-1C, was also determined by SELEX and involved G/A-rich motifs (Wu *et al*., 2016). While the U/C-rich motif associated to SR45 is similar to the binding motifs of human NXF1 and SRSF3 in the 3’UTR (3’ <u>u</u>ntranslated region) of their target RNAs (Müller-McNicoll *et al*., 2016), the ”Motif 1” is also strikingly similar to the binding motif of human NXF1 and of SRSF2, UPF3B and MLN51 (Hauer *et al*., 2016; Wegener and Müller-McNicoll, 2019). Since plant genomes lack the *TAP/NXF1* gene (Serpeloni *et al*., 2011), it is tempting to question the role of SR45 as an adaptor for the mRNA export machinery as previously reported for human SR proteins (Müller-McNicoll *et al*., 2016). It is noteworthy that the SR45-associated ACINUS is known to bind both G/A-rich and U/C-rich motifs in humans (Rodor *et al*., 2016). Different RNA motifs were uncovered for SR34 in this study **(Supplementary Figure S11 and Supplementary Tables S12-15)**, validating our SELEX approach capable of identify motifs specific to the studied RRM domain. Previous reports showed that purine-rich motifs, and specifically GAAG repeats, are typically related to positive splicing regulators (i.e. SR proteins) and are considered as exonic splicing enhancers (ESEs) in animals and plants (Fairbrother *et al*., 2002; Pertea *et al*., 2007; Thomas *et al*., 2012; Wu *et al*., 2014). Overall, this suggests that SR45 acts, to certain extent, as a positive regulator of alternative splicing (Xing *et al*., 2015; Zhang *et al*., 2017).

### Dual roles of the SR45 RRM

Our results demonstrated that the RRM of SR45 is involved in the assembly of the ASAP and PSAP complexes, and is able to efficiently bind RNA sequences. This dual role implies that its RNA binding ability and specificity might compete with the RRM contribution to protein complex formation, as previously described in the case of SRSF1 in animals (Aubol *et al*., 2016; Wegener and Müller-McNicoll, 2019; Aubol *et al*., 2021). The EJC components, Y14 and MAGO, mediate protein-protein interactions through the β-sheet surface of the RRM, which restricts the RNA binding ability of Y14 (Lau *et al*., 2003; Shi and Xu, 2003). Although SR45 associates with EJC components and EJC peripheral factors, whether it is or not intrinsically involved in EJC-mediated molecular regulation and in underlying developmental processes still remains undetermined. It is worth noting that *acinus* and *pinin* loss-of-function mutants display phenotypes similar to a *sr45-1* mutant (i.e. dwarfism, delayed flowering, and ABA hypersensitivity) (Bi *et al*., 2021).

### Conclusion

Here, we characterized the contribution of the RS and RRM domains, and their constitutive residues, to the function of the Arabidopsis SR45 protein considering the many aspects defining a splicing factor: (i) the subcellular localization, (ii) the ability to interact with protein partners, including in multiprotein complexes, as confirmed by structural modelling, and (iii) the specificity of the binding to RNA. Our results demonstrate a dual role for the RRM of SR45 in both protein-protein and protein-RNA interactions, suggesting a possible competition between both functions. Structural analysis will be useful to substantiate these findings, notably in the presence of both a partner requiring the SR45 RRM to interact with and an RNA specifically bound by this domain. Furthermore, it would be of interest to determine whether the enrichment of the RNA motifs identified here also occurs within the pre-mRNA sequences of direct RNA targets of SR45 and whether it is correlated to molecular changes in the expression and in the splicing profiles of these targets. The latter would require the assessment of genome-wide alternative splicing events occurring in the loss-of-function mutant *sr45-1* at developmental stages displaying a higher abundance of SR45, such as in seedlings or in inflorescences as shown here by our expression profiling. The mutant variants engineered in this study could serve in complementation assays of the *sr45-1* mutant with both SR45 isoforms using native promoter-driven translational fusions. Moreover, it was shown that acetylation, methylation, SUMOylation, ubiquitylation, or even prolyl *cis-trans* isomerization can regulate the activity of animal SR proteins and other splicing factors (Slišković *et al*., 2022; Kretova *et al*., 2023). Therefore, dissecting more precisely each of the SR45 isoform domains and focusing on all types of post-translational modifications would demonstrate the importance of specific residues associated to the molecular function of SR45 and to its physiological responses in the plant.

## Supplementary data

**Supplementary Figure S1.** Scaled schemes depicting the native *SR45* gene and the engineered mutant variants.

**Supplementary Figure S2.** Toxicity and autoactivation assays for yeast vectors harboring the native *SR45* or mutant variant coding sequences.

**Supplementary Figure S3.** Reporter lines depicting the activity of the native *SR45* promoter (*pSR45:EGFP*) at different developmental stages.

**Supplementary Figure S4.** Interpretive summary about the interactions of SR45 with kinases, splicing-associated factors, and nuclear import and mRNA export factors, as observed by directed yeast-two-hybrid assays.

**Supplementary Figure S5.** Interpretive summary about the interactions of SR45 with EJC core components and peripheral factors.

**Supplementary Figure S6.** SR45 interactions with all the nineteen Arabidopsis SR proteins.

**Supplementary Figure S7.** Model of the Arabidopsis ASAP assembly.

**Supplementary Figure S8.** Model of the Arabidopsis PSAP assembly.

**Supplementary Figure S9.** The mutated RRM of SR45 does not bind to any RNA consensus.

**Supplementary Figure S10.** Trinucleotide frequency among the sequences used to identify SR45-associated RNA consensuses.

**Supplementary Figure S11.** RNA motifs binding to RRMs of SR34.

**Supplementary Tables S1-9.** List of primers used in this study, subdivided into experimental categories.

**Supplementary Tables S10-15.** List of sequences submitted to MEME to find significant consensus(es) for the native RRMs studied or to find non-significant motifs associated to mutated RRMs.

**Supplementary Data S1.** Model of the Arabidopsis ASAP assembly (as pdb file).

**Supplementary Data S2.** Model of the Arabidopsis PSAP assembly (as pdb file).

## Acknowledgments

We thank Dr. Julien Spielmann and Prof. Moreno Galleni for helpful discussions.

## Author contributions

PM conceived and directed the research. SF conducted most experiments, with contributions of MS, MJ, SDF and MV for SELEX experiments. FK performed AlphaFold computational modelling. PM, FK and SF analyzed the data. PM and MH supervised experiments. SF made the figures, with the help of FK and SDF for computational modelling. SF, FK, MH and PM wrote the manuscript, with comments of all authors.

## Data availability

All data supporting the findings of the study are available within the paper and within its supplementary materials published online, or upon request.

