## Supplementary for "The *Arabidopsis* SR45 splicing factor bridges the splicing machinery and the exon-exon junction complex"

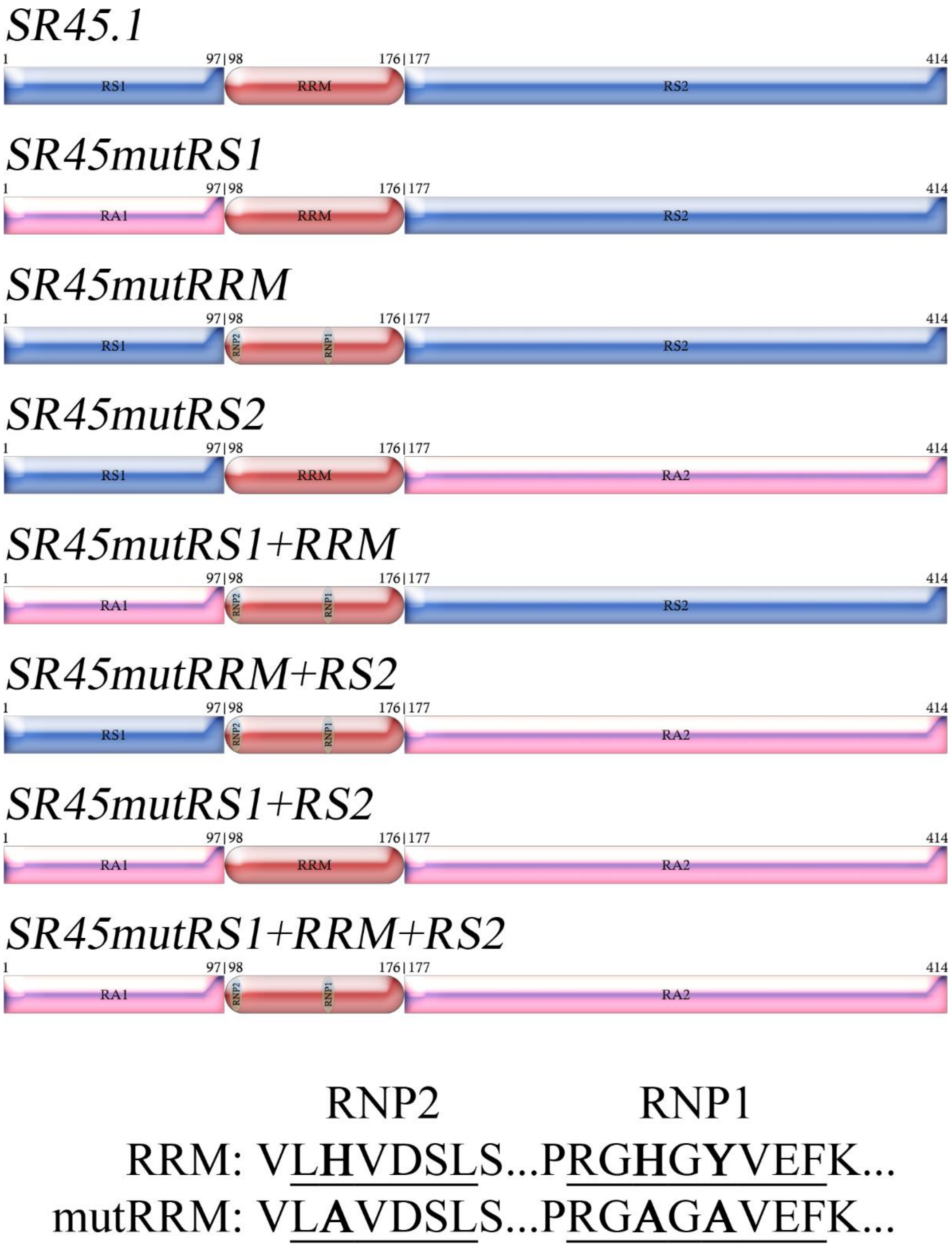


**Supplementary Figure S1.** Scaled schemes depicting the native SR45.1 protein and mutant variants. RA1 and RA2 variants correspond to RS domains in which all the serines (39 in RS1 and 49 in RS2) were substituted in alanines, respectively as shown in **Supplementary Table S6**. Critical aromatic amino acids (bold underlined) connecting the RNA in RNP2 and RNP1 motifs were substituted in alanines (Califice *et al.*, 2012; Maris *et al.*, 2005; Stankovic *et al.*, 2016). 1-414: SR45.1 protein size in amino acids.

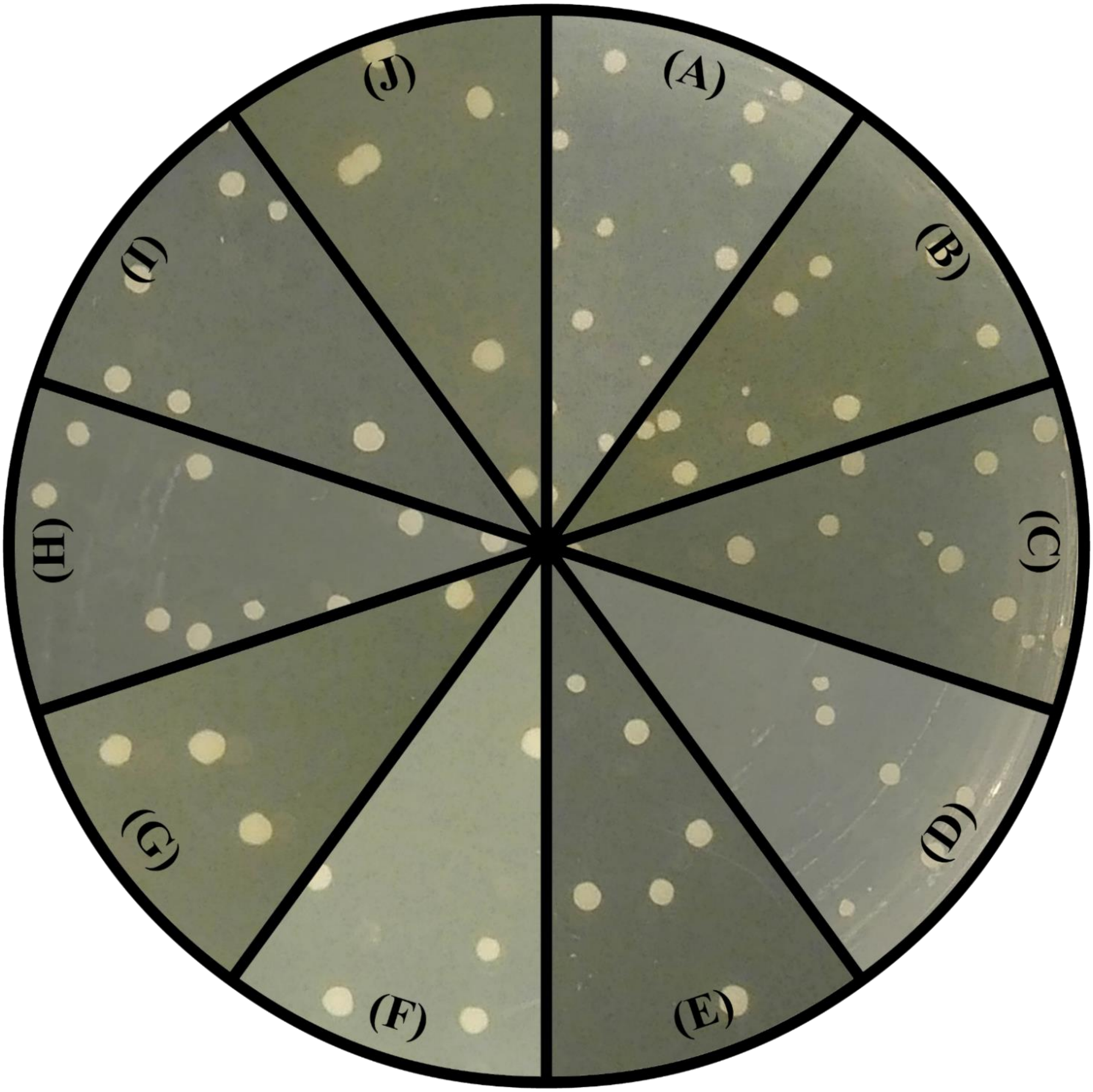


**Supplementary Figure S2.** Toxicity and autoactivation assays of the native SR45 and SR45 mutant variants on a –Trp/X-a-gal medium. Yeast transformants obtained using 100 ng of the following plasmids: **(A)** empty pGBTK7(+) vector, **(B)** pGBKT7:SR45.1, **(C)** pGBKT7:SR45.2, **(D)** pGBKT7(+):SR45mutRS1, **(E)** pGBKT7(+):SR45mutRRM, **(F)** pGBKT7(+):SR45mutRS2, **(G)** pGBKT7(+):SR45mutRS1+RS2, **(H)** pGBKT7(+):SR45mutRS1+RRM, **(I)** pGBKT7(+):SR45mutRRM+RS2, and **(J)** pGBKT7(+):SR45mutRS1+RRM+RS2. Similar results were obtained for pGADT7(+) constructs.

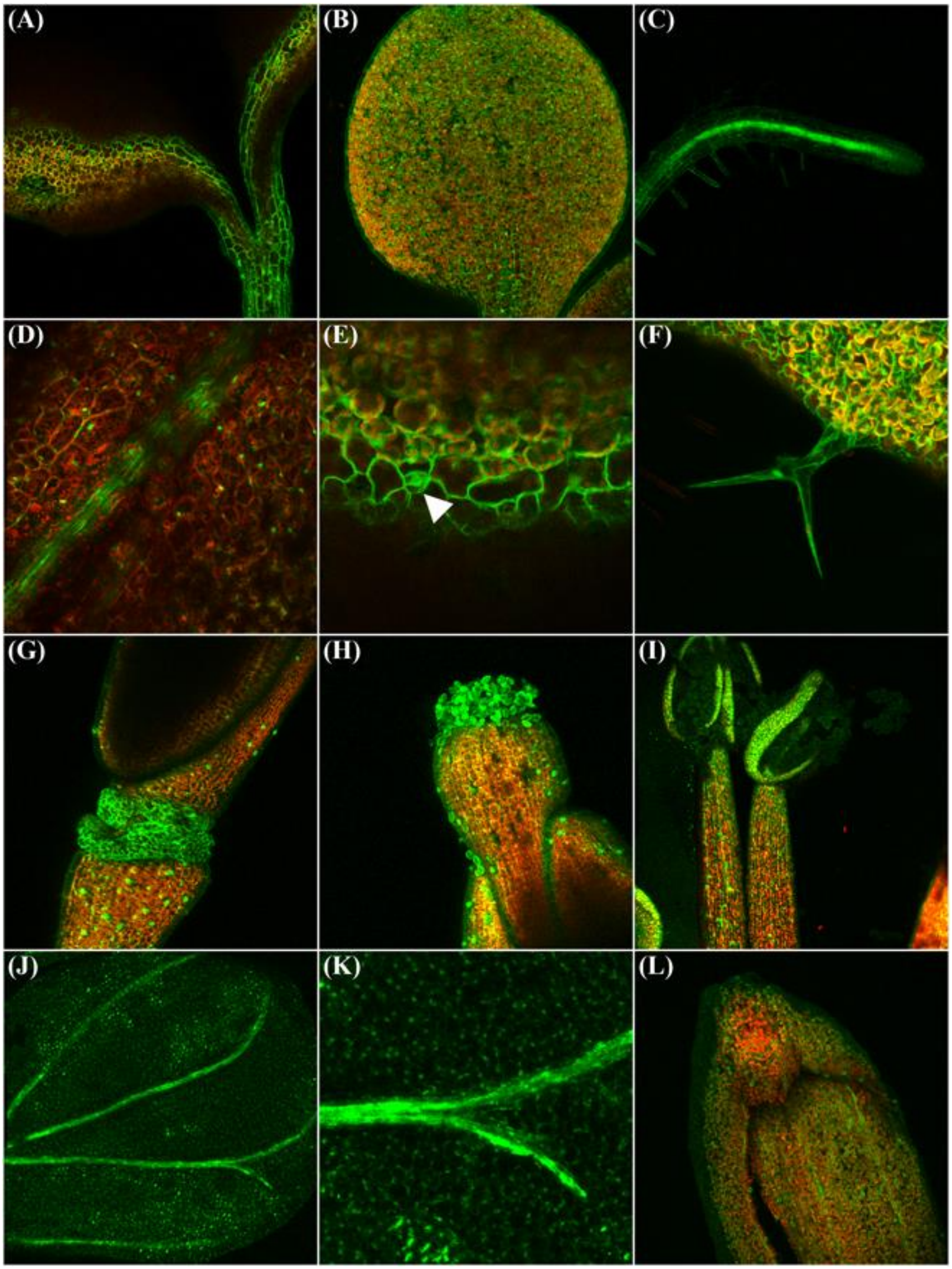


**Supplementary Figure S3.** Reporter lines expressing *pSR45:EGFP*. The promoter is active in **(A)** hypocotyl and **(A-B)** cotyledons, **(C)** two-week-old root vasculature and epidermis, **(D-F)** leaves, including epidermis and veins **(D)**, stomata [**(E)**, arrow] and trichome **(F)**, **(G-H)** gynoecium (valves, gynophore, abscission zone and receptacle) **(G)**, and stigma and style **(H)**, **(I)** androecium (anther and filament), **(J-K)** petals (epidermis and veins), and **(L)** sepal epidermis. Red signals represent chlorophyll autofluoresence. At least three independent T3 lines were generated and analyzed, depicting similar fluorescence profiles.

| | Kinases | | | | Splicing-associated factors | | | | | | | | | | | | | Nuclear imp. mRNA export | |
|---|---|---|---|---|---|---|---|---|---|---|---|---|---|---|---|---|---|---|---|
| | SRPK3a | SRPK3b | SRPK4 | AFC2 | SR34 | RSZ21 | SC35 | RS31a | RS2Z32 | SCL30 | RZ-1B | RZ-1C | PRP38 | MOS12 | Cyp59 | CypRS64 | CypRS92 | MOS14 | ALY4 |
| SR45.1 | + | + | + | + | + | + | + | + | + | + | + | + | + | + | + | + | + | + | + |
| SR45mutRS1 | − | − | − | − | + | − | − | + | ± | + | ± | − | + | − | − | − | − | ± | ± |
| SR45mutRRM | + | + | + | + | + | + | + | + | + | + | ± | ± | ± | + | − | − | − | ± | ± |
| SR45mutRS2 | − | − | + | + | + | + | − | + | + | + | + | + | + | + | − | − | − | − | + |
| SR45mutRS1+RRM | − | − | − | − | + | − | − | + | − | + | − | − | ± | − | − | − | − | − | ± |
| SR45mutRRM+RS2 | − | − | + | + | + | + | − | + | + | + | + | + | ± | + | − | − | − | − | + |
| SR45mutRS1+RS2 | − | − | − | − | − | − | − | − | − | − | − | − | + | − | − | − | − | − | ± |

| | SRPK1 | SRPK2 | AFC1 | AFC3 |
|---|---|---|---|---|
| SR45.1 | − | − | − | − |

| | pGBKT7-53 | pGBKT7-Lam |
|---|---|---|
| pGADT7-T | + | − |

| | GRP7 | GRP8 | Cyp65 | Cyp71 |
|---|---|---|---|---|
| SR45.1 | − | − | − | − |

**Supplementary Figure S4.** Interpretive summary about the interactions of SR45 with kinases, splicing-associated factors, and nuclear import and mRNA export factors, as observed by directed yeast-two-hybrid assays. Signs: +, positive interaction; -, absence of interaction; ±, weakened interaction. Companion to **Figure 3**.

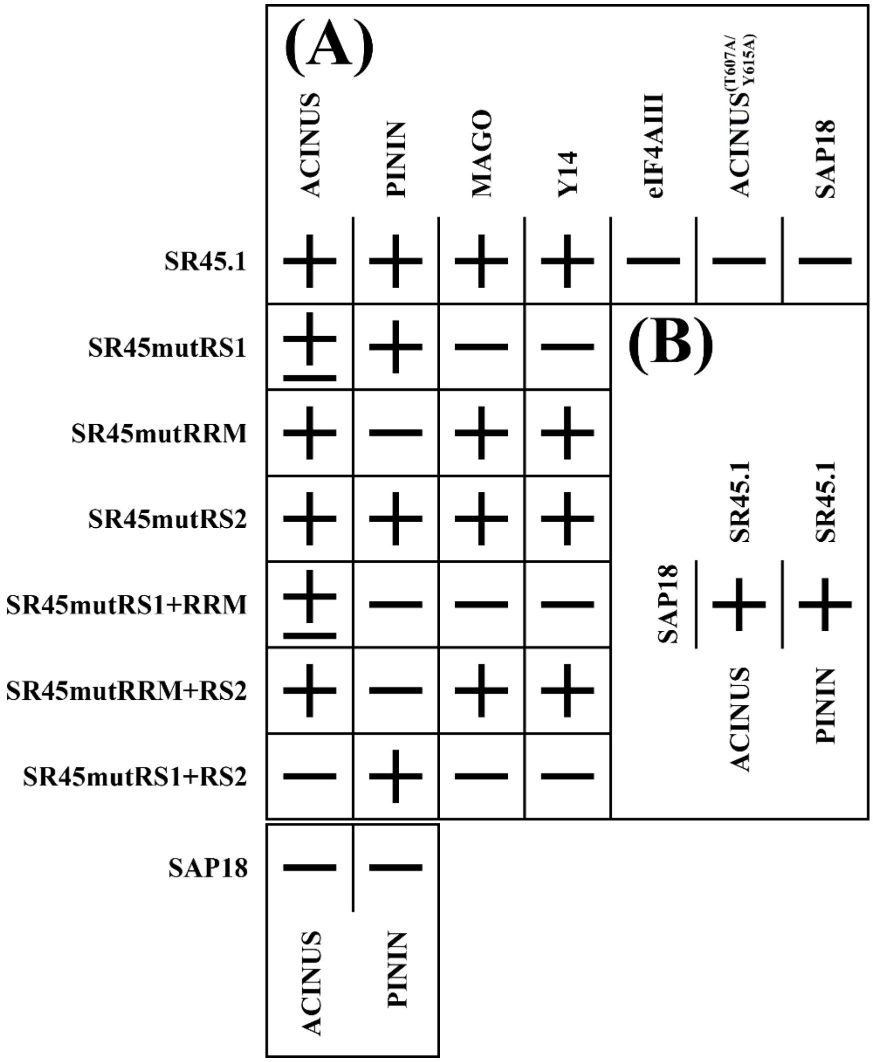


**Supplementary Figure S5.** Interpretive summary about the interactions of SR45 with EJC core components and peripheral factors. Signs: +, positive interaction; -, absence of interaction; ±, weakened interaction. Companion to **Figure 4**.

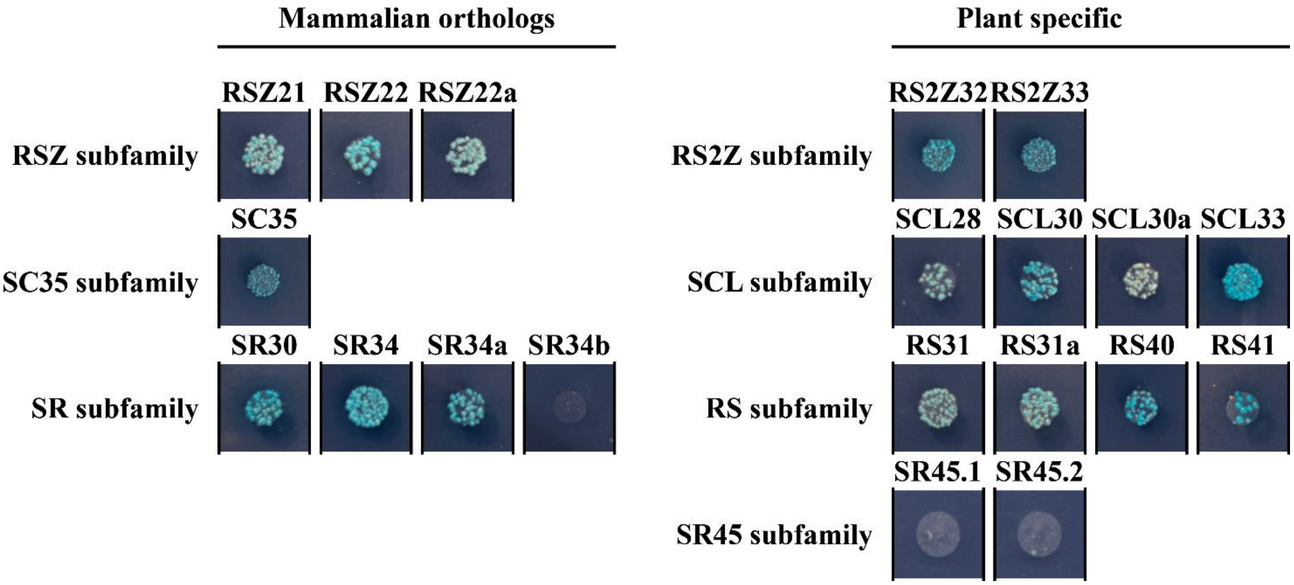


**Supplementary Figure S6.** SR45 interactions with Arabidopsis SR proteins. SR45.1 interactions with RSZ, SC35 and SR subfamilies (mammalian orthologs), and RS2Z, SCL, RS and SR45 subfamilies (plant specific). From the mated culture, dilutions to an OD600 of 0.5 were spotted on -Trp/-Leu/-His/X-a-Gal/AurA agar plates. Positive interactions were confirmed by growth and blue staining. The absence of interaction is characterized at most by a white halo of dead cells.

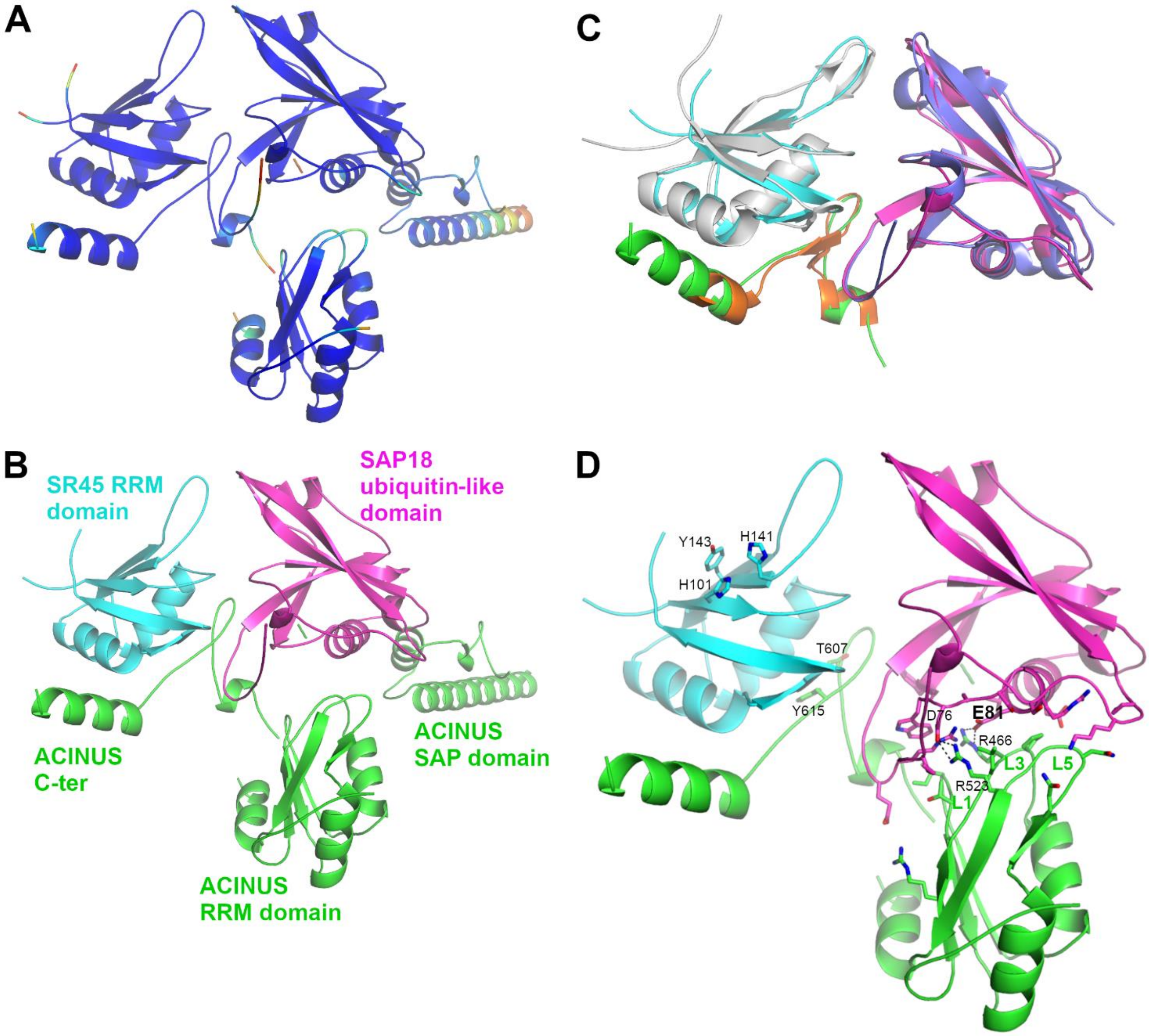


**Supplementary Figure S7.** AlphaFold Model of the *A. thaliana* ASAP complex. **(A)** Cartoon representation of the ASAP complex colored according to the pLDDT value from red (50) to yellow, cyan and blue (90). Residues with pLDDT values below 50 are considered unstructured and are not displayed. **(B)** Cartoon representation with the SR45 RRM domain in cyan, the SAP18 ubiquitin-like domain in magenta and the ACINUS SAP and RRM domains, and C-terminal end in green. **(C)** Superimposition of the *A. thaliana* ASAP complex with the same coloring scheme as **(B)** and the 4A8X ASAP structure consisting of the *H. sapiens* RNPS1 in grey, the *M. musculus* SAP18 in purple and the *D. melanogaster* ACINUS C-terminal peptide in orange (Murachelli *et al.*, 2012). **(D)** Similar to **(B)** with residues of SR45 important for RNA binding and mutated into Ala in this study in cyan sticks; T607 and Y615 from the C-terminal peptide of ACINUS important for binding to SR45 RRM domain in green sticks; residues at the interface of the RRM domain of ACINUS and the ubiquitin-like domain of SAP18 are represented as green and magenta sticks, respectively. The two salt bridges are shown as black dashed lines.

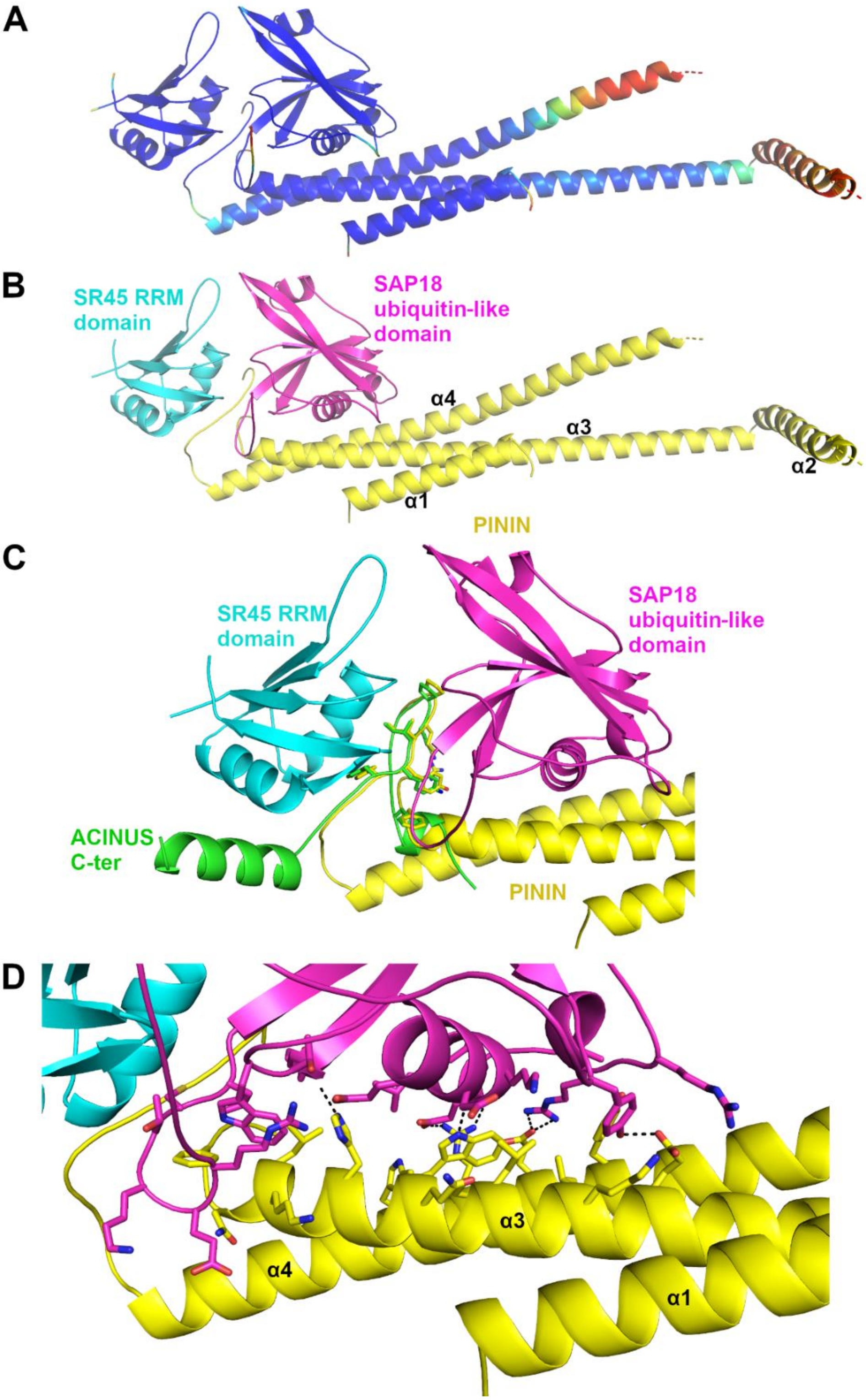


**Supplementary Figure S8.** AlphaFold Model of the *A. thaliana* PSAP complex. **(A)** Cartoon representation of the PSAP complex colored according to the pLDDT value from red (50) to yellow, cyan and blue (90). Residues with pLDDT values below 50 are considered unstructured and are not displayed. **(B)** Cartoon representation with the SR45 RRM domain in cyan, the SAP18 ubiquitin-like domain in magenta and PININ in yellow. **(C)** PSAP complex with the C-terminal peptide of ACINUS superimposed; residues conserved in ACINUS and PININ are displayed as sticks. **(D)** Interface between the ubiquitin-like domain of SAP18 and the helical region of PININ; residues within 3.5Å of the other protein are displayed as sticks. The H-bonds and salt bridges are shown as black dashed lines.

### Mutated RRM of SR45

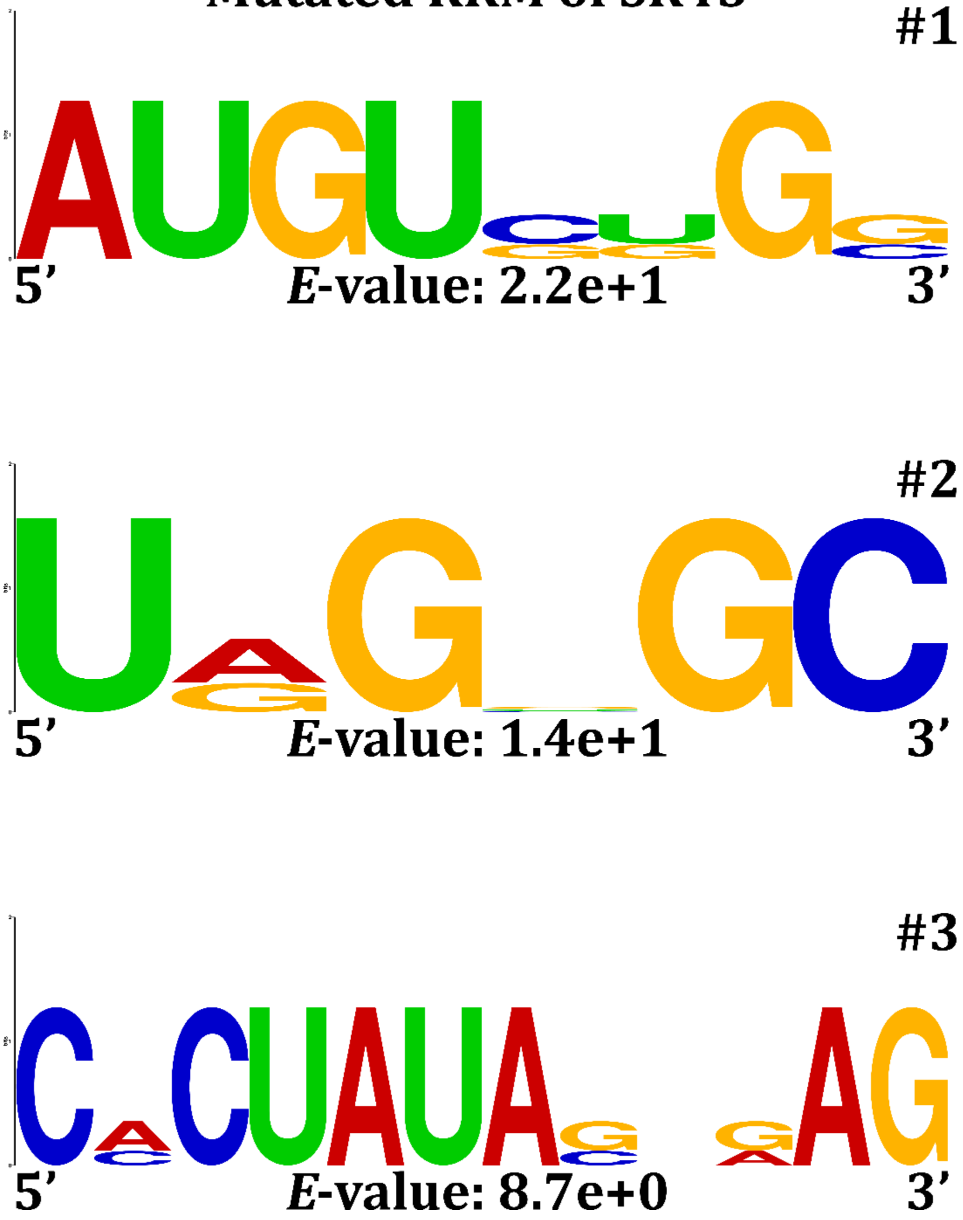


**Supplementary Figure S9.** The mutated RRM of SR45 does not bind to a specific RNA consensus. Three of the four non-significant RNA motifs (#1-3) identified through 4 rounds of SELEX selection with the mutated version of SR45 RRM are displayed. The statistical significance (*E*-value) is indicated at the bottom of each consensus. Logos were redesigned using WebLogo (Crooks *et al.*, 2004). The fourth motif (#4) (which presents the best *E*-value) is depicted in **Figure 7**. All RNA motifs were discovered using the MEME tool (version 5.4.1) with 12 sequences.

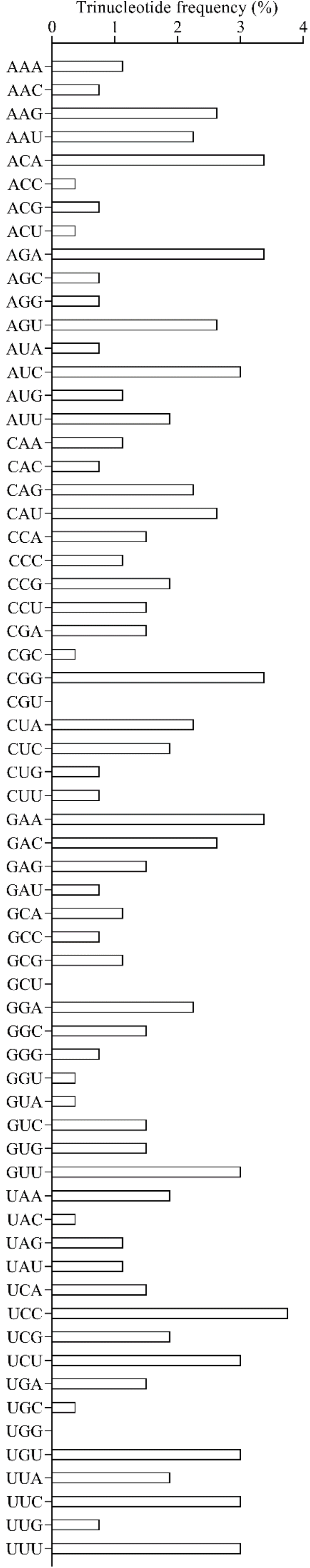


**Supplementary Figure S10.** Trinucleotide frequency among the sequences used to identify SR45-associated RNA consensuses (related to **Figure 7**).

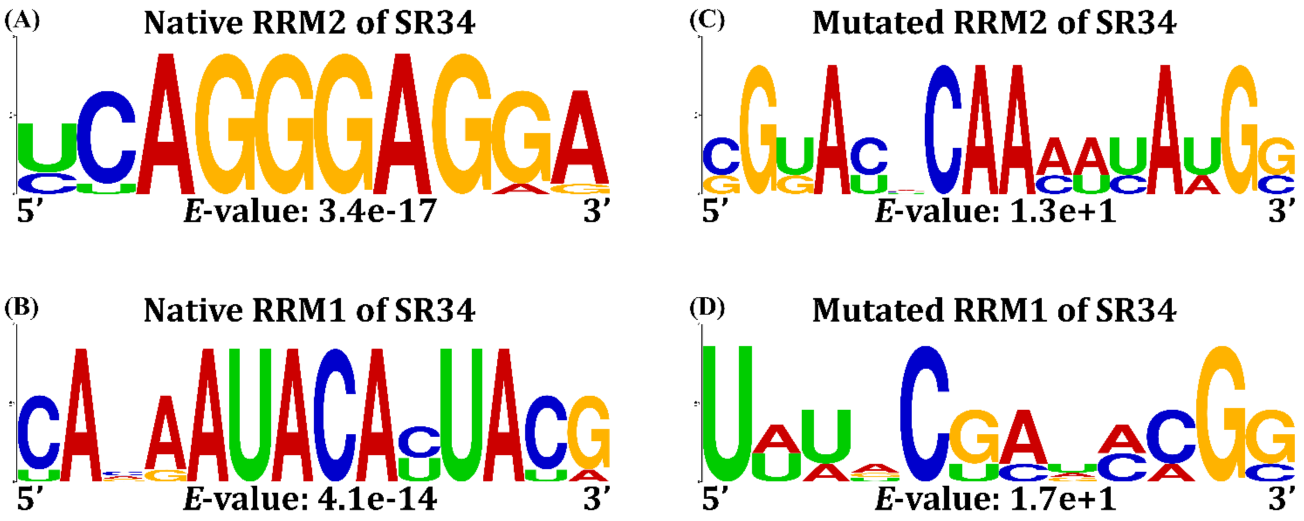


**Supplementary Figure S11.** RNA motifs binding to the RRMs of SR34. RNA motif consensuses identified through 4 rounds of SELEX selection with the RRMs of SR34 [RRM2 **(A)** or RRM1 **(B)**]. Non-significant RNA motifs identified using mutated versions of RRM2 **(C)** and RRM1 **(D)** of SR34. The statistical significance (*E*-value) is indicated at the bottom of each consensus. Logos were redesigned using WebLogo (Crooks *et al.*, 2004). All RNA motifs were discovered using the MEME tool (version 5.4.1). **(A)** The pseudo-RRM (RRM2) of SR34 recognized an invariant motif (5'-AGGGAG-3') succeeded by [G/A] and [A/G] nucleotides (*E*-value: 3.4 x $10^{-17}$), and most of the sequences are containing the AGGGAGGA motif **(Supplementary Table S12)**. This motif is in complete agreement with previously uncovered RNA sequences specifically bound by the RRM2 of the human ortholog of SR34 (SRSF1) (Cléry *et al.*, 2013; Cléry *et al.*, 2021). **(B)** The canonical RRM (RRM1) of SR34 recognized an invariant sequence (5'-AUACA-3') flanked by and enriched in CN dinucleotides ('N' for any nucleotide) (*E*-value: 4.1 x $10^{-14}$). This motif is again in agreement with a cytosine-rich motif preferentially bound by the RRM1 of SRSF1 (Cléry *et al.*, 2013; Cléry *et al.*, 2021). All sequences contained CA and CG dinucleotides **(Supplementary Table S13)**. A higher occurrence of CA dinucleotides preceding the invariant 5'-AUACA-3' was observed, predominantly in a single copy or sporadically in tandem. The CU dinucleotide was directly located after the invariant sequence in most of the cases and was usually present in a single copy. Similarly, the CG dinucleotide was mainly present in a single copy but was frequently spaced out by three nucleotides from the invariant CA dinucleotide. The least represented dinucleotide was CC.

**Supplementary Data S1.** AlphaFold Model of the Arabidopsis ASAP complex (as pdb file). This model depicts the involvement of the RRM of SR45, the ubiquitin-like domain of SAP18 and the the C-terminal peptide of ACINUS in the Arabidopsis ASAP complex assembly.

**Supplementary Data S2.** AlphaFold Model of the Arabidopsis PSAP complex (as pdb file). This model depicts the involvement of the RRM of SR45, the ubiquitin-like domain of SAP18, and both the helical region and the $^{310}$KAEPRIYYAPVKPL$^{323}$ peptide of PININ in the Arabidopsis PSAP complex assembly.

| Insertion of restriction sites in yeast plasmids | | | |
|---|---|---|---|
| **Plasmid** | **Forward (5’ – 3’)** | **Reverse (5’ – 3’)** | **Construct** |
| pGADT7 / pGBKT7 | **AATTC**CACCCGGGTGGGCATCGATACA<u>GGCGCGCC</u>GGAT<u>TTAATTAA</u>GG**G** | **GATCC**CC<u>TTAATTAA</u>ATCC<u>GGCGCGCC</u>TGTATCGATGCCCACCCGGGTG**G** | pGADT7(+) / pGBKT7(+) |

**<u>Supplementary Table S1:</u>** Primers used to insert new restrictions sites (*Asc*I and *Pac*I) in the multiple cloning sites of yeast plasmids (pGADT7 and pGBKT7). *Eco*RI and *Bam*HI sites were used for the ligation (respectively colored in orange and green).

| Target | Forward (5' – 3') | Reverse (5' – 3') | Construct | Literature |
|---|---|---|---|---|
| **SR proteins** | | | | |
| RS2Z32 | GGGAATTCATGCCTCGCTATGATGATCGC | GGGGATCCTCAAGGTGACTCACTGCCTTTAGG | pGADT7 / pGBKT7 | This study |
| RS2Z33 | GGGAATTCATGCCTCGCTATGATGATCGC | GGATCCAGGAGACTCACTTCCTCTAGG | pGADT7 / pGBKT7 | This study |
| RS31 | GGGAATTCATGAGGCCAGTGTTCGTCG | GGGGATCCTCAAGGTCTTCCTCTTGGGACT | pGADT7 | This study |
| RS31a | GGGAATTCATGAGACATGTGTACGTTGGGAATT | GGGGATCCTCAACCTCTTGCTCTTTGAATCG | pGADT7 / pGBKT7 | This study |
| RS40 | GGGAATTCATGAAGCCAGTCTTCTGTGGG | GGGGATCCTCACTCGTCAGCTGGTGGC | pGBKT7 | This study |
| RS41 | GGGAATTCATGAAGCCTGTCTTTTGCGG | GGGGATCCTCATTCCTCTGCTGGCGG | pGADT7 / pGBKT7 | This study |
| RSZ21 | GGCCCGGGTATGACGAGGGTTTATGTCGGG | GGGGATCCTCACACCCCATTGGCATATG | pGADT7 | This study |
| RSZ22 | GAATTCATGTCACGTGTGTACGTCG | GGATCCTCAGCTCCTGCTTCTGC | pGADT7 / pGBKT7 | This study |
| RSZ22a | GGGAATTCATGTCGCGTGTGTATGTTGGTAAT | GGCCCGGGTCAGCTCCGGCTTCTGC | pGADT7 / pGBKT7 | This study |
| SC35 | GGGAATTCATGTCGCACTTCGGAAGGTC | GGGGATCCTCATTCCGCAGCATAAGGAGA | pGADT7 / pGBKT7 | This study |
| SCL28 | GGGAATTCATGGCTAGAGCGAGAAGCCG | GGGGATCCTCAACGACTTAAGGATCGAGAACG | pGBKT7 | This study |
| SCL30 | GGGAATTCATGAGGAGATACAGTCCGCCTTATTA | GGCTGCAGTCATCTTGGAGATACCTCCACAGAC | pGBKT7 | This study |
| SCL30 | GGGAATTCATGAGGAGATACAGTCCGCCTTATTA | GGGAGCTCTCATCTTGGAGATACCTCCACAGAC | pGADT7 | This study |
| SCL30a | GGGAATTCATGAGAGGAAGGAGCTACACGC | GGCCCGGGTCACTGGCTTGGAGAACGG | pGADT7 / pGBKT7 | This study |
| SCL33 | GGGAATTCATGAGGGGAAGGAGCTACACTCC | GGGGATCCTCACTGGCTTGGTGAACGG | pGADT7 / pGBKT7 | This study |
| SR30 | GGGGCCATGGGGATGAGTGGGCGATTTTCTCGGTC | GGATCCACCAGATATCACAGGTGAAAC | pGADT7 / pGBKT7 | (Stankovic et al., 2016) |
| SR34 | GAATTCATGAGCAGTCGTTCGA | AGGGATCCCCTCGATGGACTC | pGADT7 / pGBKT7 | (Stankovic et al., 2016) |
| SR34a | GGGCCATGGGGATGAGTGGGCGATTTTCTCGGTC | AGGGATCCCACACTGCCTTCGC | pGADT7 / pGBKT7 | (Stankovic et al., 2016) |
| SR34b | GAATTCATGAGCAGCCGTTCG | GGATCCTATCAATGCGATCCAATG | pGBKT7 | |
| SR45.1 / SR45.2 | CGGGATCCGGATGGCGAAACCAAGTCGTGGC | CCGAGCTCTTAAGTTTTACGAGGTGGAGGTGGTGG | pGADT7 | (Stankovic et al., 2016) |
| SR45.1 / SR45.2 | CGGGATCCGGATGGCGAAACCAAGTCGTGGC | GGCTGCAGAGTTTTACGAGGTGGAGG | pGBKT7 | (Stankovic et al., 2016) |
| SR45mut(RRM+)RS2 | GGGGCGCGCCATGGCGAAACCAAGTCGTG | GGGGATCCAGTTTTACGAGGTGGAGGTGGTGGTGG | pGADT7(+) / pGBKT7(+) | This study |
| SR45mutRS1(+RRM+RS2) | GGGGCGCGCCATGGCGAAACCAGCTCGT | GGGGATCCAGTTTTACGAGGTGGAGGTGGTGGTGG | pGADT7(+) / pGBKT7(+) | This study |

| | | | | |
|---|---|---|---|---|
| **hnRNP-like** | | | | |
| GRP7 | GGGAATTCATGGCGTCCGGTGATGTTG | GGGGATCCTTACCATCCTCCACCACCACC | pGADT7 | This study |
| GRP8 | GGGAATTCATGTCTGAAGTTGAGTACCGGTGC | GGGGATCCTTACCAGCCGCCACCAC | pGADT7 / pGBKT7 | This study |
| RZ-1B | CCGAATTCATGAAAGATAGAGAAAACGATGGAAATC | GGGGATCCCTACCAACGTTCATATGATGAAGGTC | pGBKT7 | This study |
| RZ-1C | CCGAATTCATGGCTGCAAAAGAAGGTAGTAGG | GGGGATCCTTAATAACGGTCAAAAGTGGACG | pGBKT7 | This study |
| **PRPs** | | | | |
| PRP38 | GGGGATCCTTATGGCAAACAGAACAGATCCG | GGCTCGAGGTCCCTGAGGGGTTTCATTCC | pGADT7 | This study |
| **Kinases** | | | | |
| AFC1 | GGCCCGGGTATGCAAAGCAGTGTGTATCGTGATAA | GGGGATCCTTAGTTCTTTTGGTTGTATAAAAATGGATGAG | pGADT7 | This study |
| AFC2 | GGCATATGATGGAGATGGAGCGTGTGC | GGGGATCCCTATCTTCTCCTTGCGAAAAACG | pGADT7 / pGBKT7 | This study |
| AFC3 | GGGAATTCATGATAGCTAACGGATTCGAGAGTATG | GGGGATCCTCAACTTGAGCTCTTAAAGAAAGGATG | pGADT7 | This study |
| SRPK1 | GGGAATTCATGTCTTGTTCATCTTCTTCCGGAT | GGGGATCCTCACCCTTTGATATGCAAGTTGTTC | pGADT7 | This study |
| SRPK2 | GGGAATTCATGTCGTGTTCATCCTCATCTGG | GGCCCGGGTCAAGAACATGAACCTTTGATCTGC | pGADT7 | This study |
| SRPK3a | GGGAATTCATGGCGGATGAGAAGAACGG | GGGGATCCTTAAGTACGAAGATCACGAGCTGATTG | pGADT7 / pGBKT7 | This study |
| SRPK3b/SRPK5 | GGGAATTCATGGCGGAGGACAAAAACAAC | GGGGATCCTCACTTCGGTTCAGAGACATCAATAG | pGADT7 | This study |
| SRPK4 | GGGAATTCATGGAGGCGGAGAAGTGGAACAG | CGGGATCCAATTGCTAGCTTAAGAGTGGAGGAGCTT | pGADT7 / pGBKT7 | (Stankovic et al., 2016) |
| **Cyclophilines** | | | | |
| Cyp59 | GGGAATTCATGTCAGTTCTTATTGTGACGAGCC | GGGGATCCTCATCTATCCCTTCTCTCATGTCTAGCT | pGADT7 | This study |
| CypRS64 | CGGAATTCATGACTAAAAAGAAGAATCCTAATGTTTT | TATAGAGCTCTCAATCCGCATAGCTAACCAG | pGADT7 | (Stankovic et al., 2016) |
| Cyp65 | GGGAATTCATGGGGAAGAAACAACACAGCA | GGGGATCCTTACCAGCTAGAGAAATCTTTAAACCCTG | pGADT7 | This study |
| Cyp71 | GGGAATTCATGGAGGAAGAATCTAAGAATGGCG | GGCCCGGGCTAAGATTTCGGAACGGTGACATTAA | pGADT7 | This study |
| CypRS92 | GGGAATTCATGGCAAAAAAGAAGAATCCACAGG | GGCCCGGGTTAATCATAGGCTACTAATCCCTTTTTCCC | pGADT7 | This study |
| **EJC(-related)** | | | | |
| ACINUS(RRM) | CCGAATTCATGCTTCCTGCTAATGATCAAGAAGC | CCGGATCCTCACTTGTTATTATTCGCTGCAAGTTTAG | pGADT7 / pGBKT7 | This study |
| ACINUS(RRM) (T607A/Y615A) | CCGAATTCATGCTTCCTGCTAATGATCAAGAAGC | CCGGATCCTCACTTGTTATTATTCGCTGCAAGTTTAG | pGADT7 | This study |
| ALY4 | CCGAATTCATGTCTGGAGCATTGAATATGACTCTTG | CCGGATCCTCAAGAGGTGTTCATGGCATCAGC | pGADT7 / pGBKT7 | This study |
| eIF4A3 | CCGAATTCATGGCGACAGCGAATCCTGG | CCGGATCCTCAGATAAGATCAGCTACATTCATTGGC | pGADT7 / pGBKT7 | This study |

| | | | | |
|---|---|---|---|---|
| MAGO | GGGAATTCATGGCCGCGGAAGAAGC | GGGGATCCCTAGATAGGCTTGATTTTGAAGTGCAG | pGADT7 / pGBKT7 | This study |
| PININ | GGCATATGATGGGAGACACCGCCTTG | CCGAATTCTTAGAGAACCTCATGTTTAATATCTTCC | pGADT7 / pGBKT7 | This study |
| SAP18 | CCGAATTCATGGCTGAAGCAGCGAGAAGAC | CCGGATCCTCAGTAAATTGCCACATCCAGATAATC | pGBKT7 | This study |
| Y14 | CCGAATTCATGGCGAACATAGAATCAGAAGCAGTC | CCGGATCCTCAGTAACGTCTTCTCGGACTTCTTG | pGADT7 / pGBKT7 | This study |
| **MOSes** | | | | |
| MOS12 | GGCATATGATGATTTACACTGCTATCGACAATTTTTAC | GGGGATCCTTAATGGTGCCTACGACGGTCT | pGBKT7 | This study |
| MOS14 | GGCCCGGGTATGGAGCATCAGAACGCGG | GGGGATCCTGATACAGGAGCAGTAACCAGATTCA | pGADT7 | This study |
| **Trimeric complex** | | | | |
| SAP18 | CCGGATCC**GG**ATGGCTGAAGCAGCGAGAAGAC | GGCTGCAGTCAGTAAATTGCCACATCCAGATAATC | pBridge:SAP18:SR45.1 | This study |
| SR45 | GGGCGGCCGC**C**ATGGCGAAACCAAGTCGTGGC | GGGGATCCAGTTTTACGAGGTGGAGGTGGTGGTGG | pBridge:SR45.1 | This study |

**Supplementary Table S2:** Primers used to amplify coding sequences of potential interactors used in directed yeast two- and three-hybrid assays. Restriction sites used are underlined. Bold blue bases are used to preserve the open reading frame in final construction. Cells colored in light green or light orange represent, respectively, interactors identify through yeast two-hybrid screens using either a commercially available Arabidopsis cDNA library (Mate and Plate Library-Universal Arabidopsis, Clontech) or a custom cDNA library (Make Your Own "Mate & Plate" Library System, Clontech).

| Expression profiling and protein localization | | | |
|---|---|---|---|
| **Target** | **Forward (5' – 3')** | **Reverse (5' – 3')** | **Construct**<br>***Remark*** |
| pSR45 | GGCCTGCAGGATTTCTGCTCCAAAATTCTATATAAGGG | GGGGTACCGGGCGAGAGATATCGAGAAAATC | pMDC100:pSR45:SR45:EGFP:tNOS |
| SR45.1 | GGGGCGCGCCATGGCGAAACCAAGTCGTG | GGTTAATTAAAGTTTTACGAGGTGGAGGTGGTG | pMDC100:pSR45:SR45:EGFP:tNOS |
| tNOS | GGGAGCTCGAATTTCCCCGATC | GGGAGCTCAGTAACATAGATGACACCGCGC | pMDC100:pSR45:SR45:EGFP:tNOS |
| EGFP | GGTTAATTAA**T**ATGGTGAGCAAGGGCGAG | GGTTAATTAATTACTTGTACAGCTCGTCCATGCC | pMDC100:pSR45:SR45:EGFP:tNOS |
| mCherry | GGTTAATTAA**T**ATGGTGAGCAAGGGCGAG | GGTTAATTAATCATCTAGAGAATTCCTTGTACAGCTCG | pMDC100:pSR45:SR45:EGFP:tNOS |
| EGFP | CGGGCGCGCCATGGTGAGCAAGGGCGAGGAG | CCGAGCTCTTACTTGTACAGCTCGTCCATGC | pMDC32:pSR45:EGFP |
| AFC1 | GGGGCGCGCCATGCAAAGCAGTGTGTATCGTGATAA | GGTTAATTAATTAGTTCTTTTGGTTGTATAAAAATGGATGAG | pMDC32:2X35S:EGFP:AFC1 |
| AFC2 | GGGGCGCGCCATGGAGATGGAGCGTGTGC | GGTTAATTAACTATCTTCTCCTTGCGAAAAACG | pMDC32:2X35S:EGFP:AFC2 |
| AFC3 | GGGGCGCGCCATGATAGCTAACGGATTCGAGAGTATG | GGTTAATTAATCAACTTGAGCTCTTAAAGAAAGGATG | pMDC32:2X35S:EGFP:AFC3 |
| SRPK1 | GGGGCGCGCCATGTCTTGTTCATCTTCTTCCGGAT | GGTTAATTAATCACCCTTTGATATGCAAGTTGTTC | pMDC32:2X35S:EGFP:SRPK1 |
| SRPK2 | GGGGCGCGCCATGTCGTGTTCATCCTCATCTGG | GGTTAATTAATCAAGAACATGAACCTTTGATCTGC | pMDC32:2X35S:EGFP:SRPK2 |
| SRPK3a | GGGGCGCGCCATGGCGGATGAGAAGAACGG | GGTTAATTAATTAAGTACGAAGATCACGAGCTGATTG | pMDC32:2X35S:EGFP:SRPK3a |
| SRPK3b | GGGGCGCGCCATGGCGGAGGACAAAAACAAC | GGTTAATTAATCACTTCGGTTCAGAGACATCAATAG | pMDC32:2X35S:EGFP:SRPK3b |
| SRPK4 | GGGGCGCGCCATGGAGGCGGAGAAGTGGAACAG | GGTTAATTAAAATTGCTAGCTTAAGAGTGGAGGAGCTT | pMDC32:2X35S:EGFP:SRPK4<br>*No fluorescence detectable* |
| SRPK4 | GGGGATCCATGGAGGCGGAGAAGTGGAACAG | GGGGTACCAATTGCTAGCTTAAGAGTGGAGGAGCTT | pBI:35S:SRPK4:EGFP<br>*Fluorescence detectable* |
| ACINUS | GGGGCGCGCCATGTCGTCATCGCCTTTTCC | GGTTAATTAATCACTTGTTATTATTCGCTGCAAGTTTA | pMDC32:2X35S:EGFP:ACINUS |
| SAP18 | GGGGCGCGCCATGGCTGAAGCAGCGAGAAGAC | GGTTAATTAACTAGTAAATTGCCACATCCAGATAATC | pMDC32:2X35S:EGFP:SAP18 |
| PININ | GGGGCGCGCCATGGGAGACACCGCCTTG | GGTTAATTAATTAGAGAACCTCATGTTTAATATCTTCC | pMDC32:2X35S:EGFP:PININ |
| MAGO | GGGGCGCGCCATGGCCGCGGAAGAAGC | GGTTAATTAACTAGATAGGCTTGATTTTGAAGTGCAG | pMDC32:2X35S:EGFP:MAGO |
| Y14 | GGGGCGCGCCATGGCGAACATAGAATCAGAAGCAGTC | GGTTAATTAATCAGTAACGTCTTCTCGGACTTCTTG | pMDC32:2X35S:EGFP:Y14 |

**Supplementary Table S3:** Primers used for constructs for expression profiling and protein localization *in planta*. Restriction sites used are underlined. Bold blue bases are used to ensure the open reading frame in final construction.

| PCR-based point mutagenesis | | | |
|---|---|---|---|
| **Gene** | **Forward (5’ – 3’)** | **Reverse (5’ – 3’)** | **Mutant** |
| **SR45.1** | CTC**TTG**TTCTCGCTGTTGATTCTC | GAGAATCAACAGCGAGAA**CAA**GAG | SR45mutRNP2 |
| **SR45.1** | GAGGAG**CTG**GTG**CTG**TTGAG | CTCAA**CAG**CAC**CAG**CTCCTC | SR45mutRNP1 |
| **ACINUS(RRM)** | CTATTCAAGAAG**GCA**AAAGCGATCCCCAGGATATAT**GCC**TTACCCTTGT | ACAAGGGTAA**GGC**ATATATCCTGGGGATCGCTTT**TGC**CTTCTTGAATAG | ACINUS(RRM) (T607A/Y615A) |

**Supplementary Table S4:** Primers used for PCR-based point mutagenesis in the RRM domain of *SR45* and *ACINUS*. Critical aromatic amino acids connecting the RNA in RNP2 and RNP1 motifs were substituted in alanine (Maris *et al.*, 2005; Califice *et al.*, 2012; Stankovic *et al.*, 2016). The substituting codons are underlined in the sequence.

| Primers | Sequence (5’ – 3’) |
|---|---|
| **1** | GGGGCGCGCCATGGCGAAACCAAGTCGTG |
| **2** | GGGGATCCAGTTTTACGAGGTGGTGGAGGTGGTGG |
| **3** | AGAAAGCTGTTCAAGAAGCTCTTGTTCTCCATGTTGATTCTCTTAGC |
| **4** | GCAGCTACTTTCTGACGAGGTGGTAGTGTGAACGTTGCTTTAACA |
| **5** | AAGCAACGTTCACACTACCACCTCGTCAGAAAGTATCTTCACCC |
| **6** | AGAAAGCTGTTCAAGAAGCTCTTGTTCTCGCTGTTGATTCTCTTAGC |
| **7** | GGGGCGCGCCATGGCGAAACCAGCTCGT |
| **8** | GAATCAACATGGAGAACAAGAGCTTCTTGAACAGCTTTCTTAGCAG |
| **9** | GAATCAACAGCGAGAACAAGAGCTTCTTGAACAGCTTTCTTAGCAG |
| **10** | GAATCAACAGCGAGAACAAGAGATTCTTGAACAGCTTTCTTGGAAG |
| **11** | AAGCAACGTTCACACTACCACCTCGTCAGAAAGTAGCTGCAC |

**Supplementary Table S5:** Primers used to construct *SR45* mutant versions. Restriction sites used (*Asc*I et *Bam*HI) are underlined.

| Synthetic genes | |
|---|---|
| **Mutant variant** | **Coding sequence** |
| RS1 domain to RA1 domain<br><br>(291 bp, 39 S-to-A substitutions) | ATGGCGAAACCA**GCT**CGTGGCCGTCGT**GCT**CCC**GCT**GTG**GCT**GGC**GCGGCAGCT**CGT**GCTGCTGCT**AGA**GCT**CGT**GCG**GGT**GCGGCC**CCC**GCC**AGG**GCT**<br>ATT**GCT**CGC**GCA**CGC**GCT**CGT**GCT**AGA**GCG**CTC**GCTGCAGCTGCAGCT**CCT**GCG**CGA**GCC**GTC**GCTGCT**GGG**GCT**CGC**GCT**CCTCCACGTCGTGGAAAA<br>**GCT**CCTGCTGGACCTGCTAGGCGAGGTCGC**GCT**CCTCCTCCTCCACCA**GCT**AAAGGAGCA**GCTGCA**CCT**GCT**AAGAAAGCTGTTCAAGAA**GCT** |
| RS2 domain to RA2 domain<br><br>(717 bp, 49 S-to-A substitutions) | CCTCGTCAGAAAGTA**GCTGCA**CCCCCTAAACCTGTC**GCA**GCTGCACCAAAAAGAGATGCTCCAAAA**GCT**GATAATGCTGCTGCTGACGCTGAGAAAGAT<br>GGTGGTCCCAGGCGCCCAAGAGAGACA**GCT**CCTCAACGGAAAACAGGGCTT**GCT**CCGAGAAGGCGA**GCA**CCTCTACCTAGGAGAGGCTTA**GCA**CCTAGA<br>AGGCGA**GCA**CCTGAT**GCT**CCCCATCGCCGGAGGCCGGGC**GCT**CCTATCCGCCGTCGCGGTGATACACCTCCTAGACGTAGGCCAGCA**GCA**CCA**GCT**AGA<br>GGCCGT**GCT**CCA**GCTGCT**CCCCCTCCAAGACGATATAGA**GCT**CCTCCAAGGGGC**GCC**CCTAGAAGAATTCGTGGC**GCT**CCTGTTCGAAGAAGG**GCT**CCT<br>CTTCCTCTAAGACGAAGG**GCT**CCACCTCCAAGGAGACTACGC**GCT**CCTCCCAGAAGA**GCT**CCAATCCGTAGGCGT**GCT**CGA**GCT**CCAATTCGTAGGCCT<br>GGTCGC**GCT**CGT**GCAGCTGCT**ATC**GCT**CCCCGCAAGGGACGAGGTCCAGCTGGGAGACGTGGGAGG**GCAGCTGCT**TAC**GCTGCTGCA**CCA**GCT**CCCAGA<br>AGGATTCCTAGGAAGATT**GCA**AGG**GCC**CGC**GCT**CCTAAGAGGCCACTGAGAGGAAAAAGA**GCTGCCGCT**AAC**GCTGCTGCTGCTGCTGCA**CCGCCACCA<br>CCACCTCCACCTCGTAAAACTTAA |

**Supplementary Table S6:** Synthetic genes provided by Eurofins Genomics where serine codons within RS1 or RS2 of *SR45.1* gene were substituted with alanine codons (bold and underlined).

| BiFC and TriFC | | | |
|---|---|---|---|
| **Target** | **Forward (5' – 3')** | **Reverse (5' – 3')** | **Construct** |
| ACINUS(RRM) | CCGGATCCATGCTTCCTGCTAATGATCAAGAAGC | CCGGTACCCTTGTTATTATTCGCTGCAAGTTTAG | pBI121:35S:ACINUS:$^{N}$YFP |
| SR34 | CCGGATCC**GG**ATGAGCAGTCGTTCGAGTAGAACCG | GGGGTACCCCTCGATGGACTCCTAGTGTGGATAG | pBI121:35S:SR34:$^{N}$YFP |
| PRP38 | GGGGATCC**TT**ATGGCAAACAGAACAGATCCG | GGGGTACCGTCCCTGAGGGGTTTCATTCC | pBI121:35S:PRP38:$^{N}$YFP |
| ACINUS(RRM) | GGGGCGCGCCATGCTTCCTGCTAATGATCAAGAAGCTG | GGTTAATTAATCACTTGTTATTATTCGCTGCAAGTTTA | pMDC32:2X35S:ACINUS(RRM) |
| PININ | GGGGCGCGCCATGGGAGACACCGCCTTG | GGTTAATTAATTAGAGAACCTCATGTTTAATATCTTCC | pMDC32:2X35S:PININ |
| PININ | GGGGCGCGCCATGGGAGACACCGCCTTG | GGTTAATTAATTAGAGAACCTCATGTTTAATATCTTCC | pBI121:35S:PININ:$^{N}$YFP |
| SR45.1 | GGCGGGATCC**GG**ATGGCGAAACCAAGTCGTGGCCG | CCGGTACCAGTTTTACGAGGTGGAGGTGG | pBI121:35S:SR45:$^{C}$YFP |
| SR45mutRS1 and SR45mutRS1+RS2 | GGGGATCC**GG**ATGGCGAAACCAGCTCGT | CCGGTACCAGTTTTACGAGGTGGAGGTGG | pBI121:35S:SR45mutRS1:$^{C}$YFP or pBI121:35S:SR45mutRS1+RS2:$^{C}$YFP |
| pBI121 | GATCCGGCGCGCCCGGGCTCGAGACTAGTTAATTAATGGTAC | C**A**TTAATTAACTAGTCTCGAGCCCGGGCGCGCCG | pBI121:MCS (containing *Asc*I and *Pac*I) |

**Supplementary Table S7:** Primers used to confirm protein-protein interactions *in planta* (BiFC and TriFC experiments). Restriction sites used are underlined. Bold blue bases are used to ensure the open reading frame in final construction.

| SELEX | | | |
|---|---|---|---|
| **Synthesis of the DNA initial library** | | | |
| **Randomized initial library** | | ***T7* promoter** | |
| TCCCGCTCGTCGTCT*NNNNNNNNNNNNNNNNNNNNNNNNN*CCGCATCGTCCTCCCT | | GAAATTAATACGACTCACTATAGGGAGGACGATGCGG | |
| **Recombinant gene construction** | | | |
| **Target** | **Forward (5' – 3')** | **Reverse (5' – 3')** | **Construct** |
| SR34 (native & mutated RRM1) | TATAGGATCCATGAGCAGTCGTTCGAGTAGAACCG | TATAGGAATTCAACCACGGCCACCACCATTG | pGEX6P1 |
| SR34 (native & mutated RRM2) | TATAGGATCCATGGGTGGTCGCGGCCGTGGTG | TATAGGAATTCATGAATCATATTCTCTAACCCGG | pGEX6P1 |
| SR45 (native RRM) | GGGGATCCATGCTTGTTCTCCATGTTGATTCTCTTAGC | GGGAATTCTTATGGTAGTGTGAACGTTGCTTTAA | pGEX6P1 |
| SR45 (mutated RRM) | GGGGATCCATGCTTGTTCTCGCTGTTGATTCTCTTAGC | GGGAATTCTTATGGTAGTGTGAACGTTGCTTTAA | pGEX6P1 |

**Supplementary Table S8:** Primers used for SELEX experiments (De Franco *et al.*, 2019). Red bases correspond to the *T7* promoter. Restriction sites used are underlined.

| qRT-PCR | | | | |
|---|---|---|---|---|
| **Gene ID** | **Symbol** | **Forward (5’ – 3’)** | **Reverse (5’ – 3’)** | **Reference** |
| AT1G16610 | SR45.1<br>SR45.2 | CTCCAGGTCTATTTCCCGCTCA | TCCAGCAGGACTTTTTCCACG | This study |
| AT1G58050 | AT1G58050 | CCATTCTACTTTTTGGCGGCT | TCAATGGTAACTGATCCACTCTGATG | (Rausin *et al.*, 2010) |

**Supplementary Table S9:** Primers used for mRNA levels analysis (qRT-PCR).

| Native RRM of SR45 | | | | |
|---|---|---|---|---|
| **#** | **Sequence** | **Length (bp)** | **Purine (%)** | **Pyrimidine (%)** |
| **1** | UCCAGAAGUUUCCGGAAGUUCUC | 23 | 43.48 | 56.52 |
| **2** | AUUCCGGAAGUUCCCUCUAACCAUA | 25 | 40.00 | 60.00 |
| **3** | UUGAUCCGGACAUCUGCAGCACA | 23 | 47.83 | 52.17 |
| **4** | UAGUGUCUCGGCCGAGACAUUACA | 24 | 50.00 | 50.00 |
| **5** | UCCUUUGAAGGAGACAUUAUCCUA | 24 | 45.83 | 54.17 |
| **6** | UGUGUGUUCUAGAAGUUUUCUGUUUU | 26 | 34.62 | 65.38 |
| **7** | CUUAAUCGCGAUCCAGAAAUGUCAA | 25 | 52.00 | 48.00 |
| **8** | CUCAAAGGCGGCGGGGACAAACGA | 24 | 70.83 | 29.17 |
| **9** | UGUAAUGUCUCGGCCGAGACACUA | 24 | 50.00 | 50.00 |
| **10** | ACGGUCCAGACAUUUAUUUAAUCG | 24 | 45.83 | 54.17 |
| **11** | UCUAGAAGUUCCCCUAAUGAAUAU | 24 | 45.83 | 54.17 |
| **12** | GUUCAUCGGAGCAGACAUCAGUGAA | 25 | 60.00 | 40.00 |

**Supplementary Table S10:** List of sequences submitted to MEME to find two significant consensuses for the native RRM of SR45.

| Mutated RRM of SR45 | | | | |
|---|---|---|---|---|
| **#** | **1st set of sequences (Motif #1)** | **Length (bp)** | **Purine (%)** | **Pyrimidine (%)** |
| **1** | AAGAUACCCUCAAGGUAUGUGAUA | 24 | 58.33 | 41.67 |
| **2** | CGAUUAAAUAAAUGUCUGGACCGU | 24 | 54.17 | 45.83 |
| **3** | GCCAGCCGAAUGGAGACCGUAAGC | 24 | 62.50 | 37.50 |
| **4** | AAAACAGAAAACUUCUAGAACACACA | 26 | 65.38 | 34.62 |
| **5** | ACCCCACAUGGAUUCUACCUUUUA | 24 | 33.33 | 66.67 |
| **6** | UAAAAGGUAGAAUCCAUGUGGGGU | 24 | 66.67 | 33.33 |
| **7** | CGUACGGUCAAGCUCCCAACU | 21 | 42.86 | 57.14 |
| **8** | CAGAUCGGCCAUCAUGACCUAGAA | 24 | 54.17 | 45.83 |
| **9** | AAGAUGAGUCGCAAAGAAUCUCUC | 24 | 58.33 | 41.67 |
| **10** | GAGAGAUUCUUUGCGACUCAUCUU | 24 | 41.67 | 58.33 |
| **11** | UUCACUGAUGUCUGCUCCGAUGAAC | 25 | 40.00 | 60.00 |
| **12** | UUAGCACACCGCUAGUCAGCUAUA | 24 | 45.83 | 54.17 |

| Mutated RRM of SR45 | | | | |
|---|---|---|---|---|
| **#** | **2nd set of sequences (Motif #2)** | **Length (bp)** | **Purine (%)** | **Pyrimidine (%)** |
| **1** | GUUUAAGCCCUACUCGCUACAGAUC | 25 | 40.00 | 60.00 |
| **2** | GAUCUGUAGCGAGUAGGGCUUAAA | 24 | 62.50 | 37.50 |
| **3** | AGUACAUACACCUAGAAUUAGUGC | 24 | 54.17 | 45.83 |
| **4** | GCACUAAUUCUAGGUGUAUGUACUG | 25 | 48.00 | 52.00 |
| **5** | AAUUUACAACAUAGUGCCGGUAUC | 24 | 50.00 | 50.00 |
| **6** | AACACAUUACUGGAACGCCCCAUAC | 25 | 48.00 | 52.00 |
| **7** | GUAUGGGGCGUUCCAGUAAUGUGUU | 25 | 52.00 | 48.00 |
| **8** | GACGAAACAAUUUCAUACAAACAG | 24 | 62.50 | 37.50 |
| **9** | UCAUUUUCCGCUCAAAU | 17 | 29.41 | 70.59 |
| **10** | AACCAGCAAGAGUCUAUCCGGAGA | 24 | 62.50 | 37.50 |
| **11** | AUAAGUAACCUGUGCUUGGCGC | 22 | 50.00 | 50.00 |
| **12** | UCUGUAUACAGAUACAUCUAAU | 22 | 45.45 | 54.55 |

| Mutated RRM of SR45 | | | | |
|---|---|---|---|---|
| **#** | **3rd set of sequences (Motif #3)** | **Length (bp)** | **Purine (%)** | **Pyrimidine (%)** |
| **1** | CUCGUAUAUCGUGCGCCGUUACUA | 24 | 37.50 | 62.50 |
| **2** | UGAUGAAAUUGAGAAUGUAGAUG | 23 | 69.57 | 30.43 |
| **3** | GCUCCUAUACUUGUCAUAAGAAC | 23 | 43.48 | 56.52 |
| **4** | GUUCUUAUGACAAGUAUAGGAGC | 23 | 56.52 | 43.48 |
| **5** | UAUCGCCAUGGGUAUAAUCCACUA | 24 | 45.83 | 54.17 |
| **6** | CCAAUAACUAGAAGACAAAGGGGU | 24 | 70.83 | 29.17 |
| **7** | AUCCUCGCCAAUUAAGAAGACUUA | 24 | 50.00 | 50.00 |
| **8** | UAAGUCUUCUUAAUUGGCGAGGAU | 24 | 50.00 | 50.00 |
| **9** | CCAUGUGAAAUCCAUUACAGAGUA | 24 | 54.17 | 45.83 |
| **10** | CUCCACUAUACCAAGAAAACGCAG | 24 | 54.17 | 45.83 |
| **11** | GGAUUAAUACGACUCACUAUAGGGAG | 26 | 61.54 | 38.46 |
| **12** | CUCCCUAUAGUGAGUCGUAUUAAUCC | 26 | 38.46 | 61.54 |

| Mutated RRM of SR45 | | | | |
|---|---|---|---|---|
| **#** | **4th set of sequences (Motif #4)** | **Length (bp)** | **Purine (%)** | **Pyrimidine (%)** |
| **1** | ACCAUUGACGGAGACCUUAUCAAA | 24 | 54.17 | 45.83 |
| **2** | CUCAAAGGCGGCUGGGACAAACGA | 24 | 66.67 | 33.33 |
| **3** | CCGUUAUCAAGAUGGCAAUGUAGA | 24 | 58.33 | 41.67 |
| **4** | GGAUUCACAACAUAUCUAUACAAA | 24 | 54.17 | 45.83 |
| **5** | AAAAUGACUGGAUUUGGAUUCCUU | 24 | 50.00 | 50.00 |
| **6** | GGAGGGAGCUUACUAU | 16 | 62.50 | 37.50 |
| **7** | GCGCCUAGAAUACCCCGUCAUAAA | 24 | 50.00 | 50.00 |
| **8** | CUGGUACAGUAUUUACAACCUGUA | 24 | 45.83 | 54.17 |
| **9** | GACACUUUAAAUUAUAAACUAGUC | 24 | 50.00 | 50.00 |
| **10** | CCCAUCAAAUGUCCUCUACCUUGU | 24 | 29.17 | 70.83 |
| **11** | CACAAAAUGAAUUCGAUGUUUU | 22 | 50.00 | 50.00 |
| **12** | UCGUUUGUCCCCGCCGCCUUUGAG | 24 | 29.17 | 70.83 |

**Supplementary Table S11:** List of sequences (four sets of twelve sequences) submitted to MEME for the mutated RRM of SR45.

| Native RRM2 of SR34 | | | | |
|---|---|---|---|---|
| **#** | **Sequence** | **Length (bp)** | **Purine (%)** | **Pyrimidine (%)** |
| **1** | UAUUCCUAAGCCAGGGAGGA | 20 | 60.00 | 40.00 |
| **2** | CGAUGCGAUAUACCUAAGCCAGGGAGGA | 28 | 64.28 | 35.72 |
| **3** | UACCUAGUUAGUCAGGGAGGA | 21 | 61.90 | 38.10 |
| **4** | CGAUGCGACCAGGGAGGGACGCAUUAAUCCAG | 32 | 62.50 | 37.50 |
| **5** | CAACUUCAGGGAGGA | 15 | 66.67 | 33.33 |
| **6** | CAGUUUGUCAGGGAGGACAAUACG | 24 | 62.50 | 37.50 |
| **7** | UACAAUCAGGGAGGAUGUAGUCCG | 24 | 62.50 | 37.50 |
| **8** | UGAGGUUAGGGAGAACCUUUAACG | 24 | 62.50 | 37.50 |
| **9** | UGGUCACAGCACGCAGUCAGGGAGGA | 26 | 65.38 | 34.62 |
| **10** | CGAUGCGAUGGUCACAGCACGCAGUCAGGGAGGA | 34 | 64.71 | 35.29 |

**Supplementary Table S12:** List of sequences submitted to MEME to find one significant consensus for the native RRM2 of SR34.

| Native RRM1 of SR34 | | | | |
|---|---|---|---|---|
| **#** | **Sequence** | **Length (bp)** | **Purine (%)** | **Pyrimidine (%)** |
| **1** | CAGAAUACAUUACGAACGUUAGGC | 24 | 58.33 | 41.67 |
| **2** | UCACAAUACAUUACGCACCGUACA | 24 | 45.83 | 54.17 |
| **3** | CACAAUACACUACGUACGUAGCUC | 24 | 45.83 | 54.17 |
| **4** | AAGUUUACGAUACACUACAUUGU | 23 | 47.82 | 52.18 |
| **5** | CAGAAUACAUUACGCAUGUUUCAA | 24 | 50.00 | 50.00 |
| **6** | CAAAAUACACUACGCAUCAGAUGC | 24 | 54.17 | 45.83 |
| **7** | UAGUCAAAAUACACUAUGCCGAGGGAG | 27 | 62.97 | 37.03 |

**Supplementary Table S13:** List of sequences submitted to MEME to find one significant consensus for the native RRM1 of SR34. The invariant motif 5'-AUACA-3' is underlined in the sequence. Bold red bases correspond to CA dinucleotides. Bold green bases correspond to CU dinucleotides. Bold blue bases correspond to CG dinucleotides.

| Mutated RRM2 of SR34 | | | | |
|---|---|---|---|---|
| **#** | **Sequence** | **Length (bp)** | **Purine (%)** | **Pyrimidine (%)** |
| **1** | UGUACAAAUAAAUAAGGCGUCAUA | 24 | 62.50 | 37.50 |
| **2** | AGGACAUUUUAUGCGACAACCUUU | 24 | 45.83 | 54.17 |
| **3** | GUCCGGGGACACAACUUAUGG | 21 | 57.14 | 42.86 |
| **4** | AUUGGGCCCAUGUAGCCUGUCUAU | 24 | 41.67 | 58.33 |
| **5** | GACCCCCCCACUCUAGAUAAUUAC | 24 | 37.50 | 62.50 |
| **6** | UAAAUACACGUAUUCAAAACAAGG | 24 | 62.5 | 37.5 |
| **7** | CCGCCGUACCCAAAAUAUGCUUCA | 24 | 41.67 | 58.33 |
| **8** | AAUCAACCAGUCAUUUUCCCGCAAA | 24 | 62.5 | 37.5 |
| **9** | UUCCCAUGCAAUUACCCCCUCAGA | 24 | 33.33 | 66.67 |

**Supplementary Table S14:** List of sequences submitted to MEME for the mutated RRM2 of SR34.

| Mutated RRM1 of SR34 | | | | |
|---|---|---|---|---|
| **#** | **Sequence** | **Length (bp)** | **Purine (%)** | **Pyrimidine (%)** |
| **1** | CAACAACAUGCUUUUUCGAGACGG | 24 | 50.00 | 50.00 |
| **2** | AUAUCAUCAAAAAAUGUGCGGAAUA | 25 | 64.00 | 36.00 |
| **3** | AUUUACUCUCCGGGAAAAAUACAU | 24 | 50.00 | 50.00 |
| **4** | UACAUCCGUCGCCAAUCAAAGGUG | 24 | 50.00 | 50.00 |
| **5** | UAUAGACUCAUUAAACGAUAAGGC | 24 | 58.33 | 41.67 |
| **6** | CCAAAAACGAACACAUAUCAAGAA | 24 | 66.67 | 33.33 |
| **7** | UUUAUGAUACAAGCCAAUUUGUAU | 24 | 45.83 | 54.17 |
| **8** | GGACUAUAGGGAG | 13 | 76.92 | 23.08 |
| **9** | UUCAAGAACAUAUGCGAACCGCGG | 24 | 58.33 | 41.67 |

**Supplementary Table S15:** List of sequences submitted to MEME for the mutated RRM1 of SR34.
